# Type-specific dendritic integration in mouse retinal ganglion cells

**DOI:** 10.1101/753335

**Authors:** Yanli Ran, Ziwei Huang, Tom Baden, Harald Baayen, Philipp Berens, Katrin Franke, Thomas Euler

## Abstract

Neural computation relies on the integration of synaptic inputs across a neuron’s dendritic arbour. However, the fundamental rules that govern dendritic integration are far from understood. In particular, it is still unclear how cell type-specific differences in dendritic integration arise from general features of neural morphology and membrane properties. Here, retinal ganglion cells (RGCs), which relay the visual system’s first computations to the brain, represent an exquisite model. They are functionally and morphologically diverse yet defined, and they allow studying dendritic integration in a functionally relevant context. Here, we show how four morphologically distinct types of mouse RGC with shared excitatory synaptic input (transient Off alpha, transient Off mini, sustained Off, and F-mini^Off^) exhibit distinct dendritic integration rules. Using two-photon imaging of dendritic calcium signals and biophysical modelling, we demonstrate that these RGC types strongly differ in their spatio-temporal dendritic integration: In transient Off alpha cells, dendritic receptive fields displayed little spatial overlap, indicative of a dendritic arbour that is partitioned in largely isolated regions. In contrast, dendritic receptive fields in the other three RGCs overlapped greatly and were offset to the soma, suggesting strong synchronization of dendritic signals likely due to backpropagation of somatic signals. Also temporal correlation of dendritic signals varied extensively among these types, with transient Off mini cells displaying the highest correlation across their dendritic arbour. Modelling suggests that morphology alone cannot explain these differences in dendritic integration, but instead specific combinations of dendritic morphology and ion channel densities are required. Together, our results reveal how neurons exhibit distinct dendritic integration profiles tuned towards their type-specific computations in their circuits, with the interplay between morphology and ion channel complement as a key contributor.

## INTRODUCTION

Across the nervous system, the output signal of a neuron is determined by how it integrates the often thousands of synaptic inputs it receives across its dendritic arbour (Guo et al., 2014; Koch et al., 1982). However, still little is known about how dendritic integration is shaped by differences between neuron types, such as specific dendritic morphology and ion channel complement and density. To investigate type-specific dendritic integration and the key factors driving it, we here use the vertebrate retina, a model system with a clear input-output relationship that can be recorded in a dish (Ames and Nesbett, 1981). The retina decomposes the visual signal into ∼40 feature-specific parallel channels (reviewed in (Baden et al., 2018)), relayed to the brain by a matching number of retinal ganglion cell types (Baden et al., 2016; Sanes and Masland, 2015). Retinal ganglion cells (RGCs) receive their main excitatory drive from the bipolar cells (BCs), which pick up the photoreceptor signal in the outer retina. In addition, RGCs (and BCs) receive inhibitory input from amacrine cells (ACs) (reviewed in (Diamond, 2017)), completing the canonical RGC input circuit. Different RGC types differ in morphology (Bae et al., 2018; Helmstaedter et al., 2013; Sümbül et al., 2014), synaptic connectivity (Field et al., 2010; Helmstaedter et al., 2013), and expression of ion channels (Rheaume et al., 2018; Siegert et al., 2012).

To explain the emergence of diverse RGC functions, many previous studies have focused on the selective connectivity with presynaptic neurons in the inner plexiform layer (IPL) (e.g. (Helmstaedter et al., 2013; Lee et al., 2010; Yu et al., 2018)). Different RGC types arborize in specific layers of the IPL and, hence, receive synaptic inputs from distinct combinations of BC and AC types (Helmstaedter et al., 2013). This spatiotemporally heterogeneous input provides the basis of type-specific feature extraction (Roska and Werblin, 2001). In addition, RGC dendrites may themselves perform complex computations and therefore contribute to the generation of specific output channels, e.g. through their dendritic geometry, and the complement, distribution and density of passive and active ion channels (Branco and Hausser, 2010; Goldstein and Rall, 1974; Koch et al., 1982; Lai and Jan, 2006; Stuart and Spruston, 2015). So far, dendritic processing in the retina has been studied mainly in interneurons (e.g. (Chapot et al., 2017; Grimes et al., 2010; Hausselt et al., 2007; Koren et al., 2017)). Despite some theoretical work in this direction (reviewed in (Guo et al., 2014)), experimental evidence for type-specific dendritic computation and their biophysical mechanisms in RGCs remains limited and is restricted to a few specific types (i.e. direction-selective RGCs, (Dorostkar et al., 2010; Oesch et al., 2005; Sivyer and Williams, 2013); On alpha RGCs, (Schwartz et al., 2012)).

Here, we exploit the unique structure of the IPL to isolate the contributions of type-specific synaptic input profiles from intrinsic cellular mechanisms to elucidate whether RGC types sampling from a similar input space use specific dendritic integration profiles to generate functionally diverse outputs. To this end, we studied in mouse retina the dendritic integration properties of four Off RGC types that receive excitatory input from a highly overlapping set of presynaptic neurons. To record light stimulus-evoked signals across the dendritic arbour of individual RGCs, we used two-photon Ca^2+^ imaging. We found that these morphologically diverse RGC types differed strongly in their spatio-temporal dendritic integration properties. A biophysical model suggests that the differential dendritic integration in these RGC types arises from the type-specific combination of dendritic morphology and ion channel complement.

## RESULTS

### Estimating local dendritic receptive fields in single retinal ganglion cells

To study dendritic integration in different RGC types, we recorded Ca^2+^ signals in response to visual stimulation across the dendritic arbour of individual cells in the *ex-vivo*, whole-mounted mouse retina using two-photon imaging. For that, we injected individual RGCs with the fluorescent Ca^2+^ indicator dye Oregon Green BAPTA-1 (OGB-1) using sharp electrodes (Methods), resulting in completely labelled individual cells (Fig. 1a). After recording dendritic activity, the cells were 3D-reconstructed (Fig. 1b), allowing us to extract morphological parameters such as dendritic arbour area, branching order and asymmetry. To determine the cell’s dendritic stratification profile across the IPL relative to the ChAT bands, blood vessels labelled with Sulforhodamine 101 (SR101) were used as landmarks (Fig. 1a, b; Methods).

**Figure 1.**
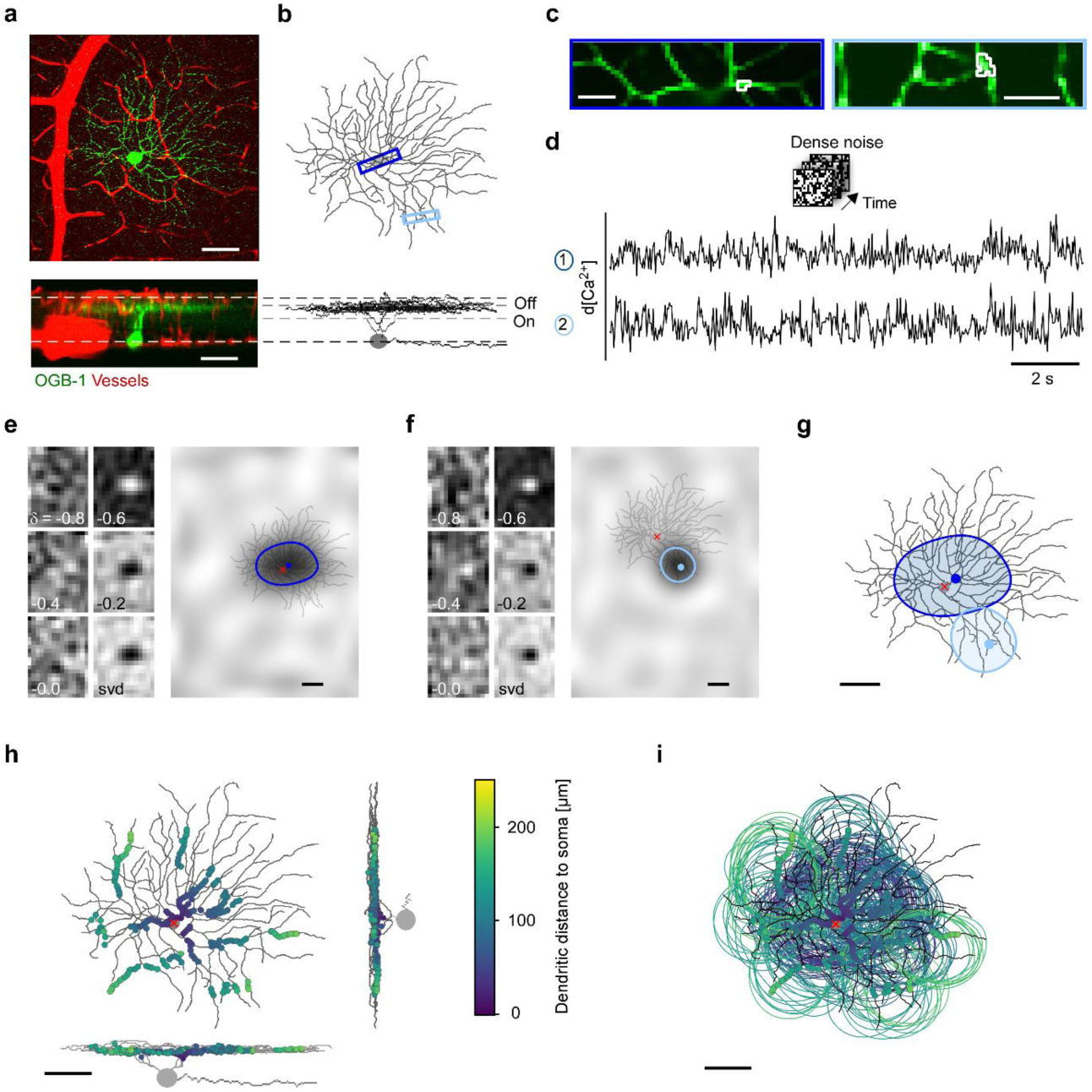
Recording dendritic receptive fields (RFs) in individual retinal ganglion cells (RGCs). **a**, Z-projection of an image stack showing an Off RGC filled with the synthetic Ca^2+^ indicator Oregon green BAPTA-1 (OGB-1; green) and blood vessels (red) in top view (top) and as side view (bottom). Dashed white lines mark blood vessels at the borders to ganglion cell layer (GCL) and inner nuclear layer (INL). **b**, Reconstructed morphology of cell from (a). Dashed grey lines between vessel plexi indicate ChAT bands. **c**, Example scan fields, as indicated by blue rectangles in (b), with exemplary region of interest (ROI; white) each. **d**, De-trended Ca^2+^ signals from ROIs in (c) during dense noise stimulation (20×15 pixels, 30 µm/pixel, 5 Hz). **e**, Spatial receptive field (RF) maps from event-triggered averaging for left ROI in (c) at different times (δ, [s]) before an event and singular value decomposition (svd; Methods) map (left). Up-sampled and smoothed RF map overlaid with the cell’s morphology (right; red crosshair indicates soma position), ROI position (blue dot) and RF contour. **f**, Like (e) but for right ROI in (c). **g**, RF contours of ROIs from (e, f) overlaid on the reconstructed cell morphology. **h**, Top- and side-view of example cell with all analysed ROIs (n=15 scan fields, n=193 of 232 ROIs passed the quality test; see Methods and Suppl. Fig. S1a, b), colour-coded by dendritic distance from soma. **i**, RF contours of ROIs from (h). Scale bars: a,b,e-i, 50 µm, c, 10 µm.

To map dendritic receptive fields (RFs) of RGCs (Fig. 1c, d), we used a binary dense noise stimulus (20×15 pixels, 30 µm/pixel) that was centred on the recording field. For each recording field (32×16 pixels @31.25 Hz), we extracted regions-of-interest (ROIs) along the dendrites using local image correlations (Supplementary Fig. S1a; Methods). Next, we registered the position and distance of each dendritic segment relative to the soma and extracted each ROI’s Ca^2+^ signal. To mitigate the effect of low signal-to-noise ratio in some dendritic recordings, we used routinely automatic smoothness determination (Methods; (Sahani and Linden, 2003)) to obtain reliable estimates of each ROI’s RF. Then, we overlaid the RF contours with the cell’s morphology (Fig. 1e-g). For each cell, we recorded different dendritic regions at various distances from the soma yielding between 40 and 232 ROIs per cell (Fig. 1h, i; Supplementary Fig. S1c). This enabled us to systematically probe dendritic integration across an RGC’s dendritic arbour and link the properties of local dendritic RFs to overall cell morphology.

### Recorded retinal ganglion cells are clustered into four morphological types

To compare dendritic integration profiles across RGC types with highly overlapping excitatory inputs, we focussed on Off RGCs that stratify close to the Off ChAT band (Fig. 2a; Supplementary Fig. S2). We recorded n=31 cells and clustered them into four morphological groups, using four morphological criteria: soma size, arbour asymmetry, arbour density difference and arbour area following Bae et al. (2018) (Fig. 2; Methods). One group likely corresponded to transient Off alpha (tOff alpha) RGCs, as indicated by a large soma and dendritic area (for statistics, see Table 1) and their characteristic stratification profile (compare to “4 ow” RGCs in the EyeWire database of reconstructed cells of the mouse retina, http://museum.eyewire.org). The second group likely represented the Off “mini” alpha transient type (tOff mini; (Baden et al., 2016)): Cells assigned to this group exhibited an IPL stratification profile very similar to tOff alpha cells, but had smaller somata and dendritic areas. The third group resembled the morphology of F-mini^Off^ cells (Rousso et al., 2016), exhibiting an IPL stratification profile peaking between the Off ChAT band and the outer IPL border and a small, highly asymmetrical dendritic arbour. Finally, the forth group displayed a similar IPL stratification profile as sustained Off alpha RGCs (“1wt” cells in (Bae et al., 2018)), but had smaller somata and arbour areas. These cells may correspond to the Off sustained (G7) RGCs identified by Baden et al (2016). Here, we refer to them as sustained Off (sOff). In the following, for simplicity, we will refer to these morphological groups as RGC types.

**Figure 2.**
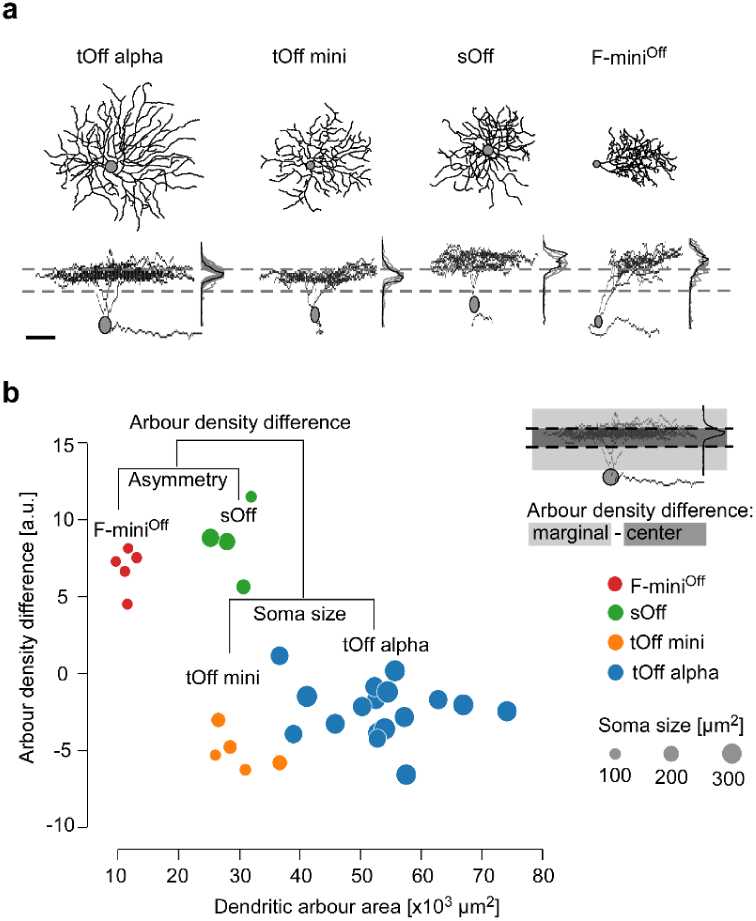
Anatomical clustering of recorded RGCs. **a**, Top- and side-views of four reconstructed Off RGCs, one of each studied type, with IPL stratification profiles as mean (black) and for all recorded cells of that type (grey). Dashed lines indicate On and Off ChAT bands. **b**, Cluster-dendrogram with the morphological parameters used in each clustering step and the resulting RGC groups: n=17 tOff alpha, n=5 tOff mini, n=4 sOff and n=5 F-mini^Off^. Colours indicate cluster (RGC type), dot diameter represents soma area. Inset: Illustration of arbour density difference measure. Scale bar: a, 50 µm.

**Table 1.** Morphological parameters describing the dendritic arbours of the clustered RGCs. For parameter definitions, see Methods.

| | n | Arbour density difference (a.u.) | Area ($10^3 \mu\text{m}^2$) | Asymmetry (a.u.) | Soma size ($\mu\text{m}^2$) |
| --- | --- | --- | --- | --- | --- |
| *tOff alpha* | 17 | $-2.41 \pm 0.44$ | $53.2 \pm 2.3$ | $44.9 \pm 6.3$ | $322.5 \pm 8.3$ |
| *tOff mini* | 5 | $-5.10 \pm 0.57$ | $29.7 \pm 1.9$ | $18.9 \pm 4.1$ | $151.5 \pm 14.8$ |
| *sOff* | 4 | $8.58 \pm 1.19$ | $29.0 \pm 1.5$ | $14.2 \pm 3.7$ | $204.0 \pm 34.1$ |
| *F-mini<sup>Off</sup>* | 5 | $6.77 \pm 0.62$ | $11.5 \pm 0.6$ | $74.8 \pm 6.2$ | $102.7 \pm 2.2$ |

### Dendritic integration profiles vary across retinal ganglion cell types

Dendrites can process incoming synaptic inputs on a local and a global scale, resulting in rather compartmentalized and synchronized dendrites, respectively (Behabadi and Mel, 2014; Bernander et al., 1991; Koch et al., 1982; Magee, 2000; Polsky et al., 2004; Williams, 2004). To investigate whether the four Off RGC types differ with respect to their integration mode, we first assessed how the RF size changed as a function of dendritic distance to the soma. In tOff alpha cells, local RF area systematically decreased as a function of ROI distance from the soma (Fig. 3a-c; *cf*. Fig. 1g, i), suggesting that signals in distal dendrites of tOff alpha cells are more isolated and local than those in proximal dendrites. This was not the case in the three remaining RGC types, where RF size remained relatively constant across different positions of the dendritic arbour (Fig. 3a-c). In fact, proximal RFs were significantly larger in tOff alpha cells than in the other RGC types (Fig. 3d; for statistical analysis, see Supplementary Information), which did not differ significantly in their RF size along their dendrite. Together, among the recorded RGC types, dendritic signals in tOff alpha cells are the least spatially synchronized, suggesting that they process dendritic input more locally than the other types.

**Figure 3.**
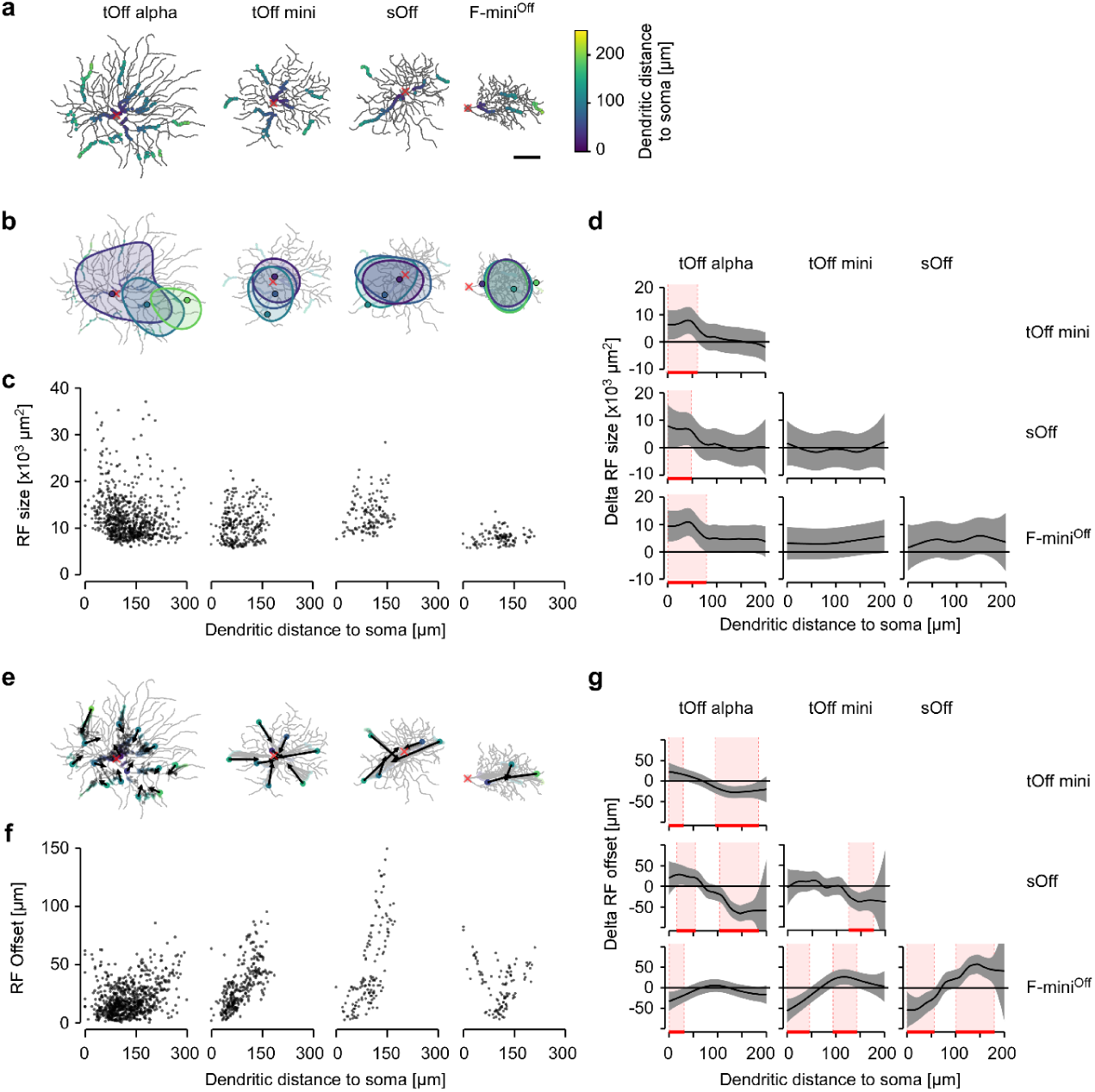
Local dendritic RF area and position varies in different RGC types. **a**, Top-views of the different reconstructed RGC types, overlaid with ROIs that passed the RF quality test. ROI colours indicate dendritic distance to soma. **b**, Cells from (a) with three ROIs of increasing distance from soma and corresponding RF contours overlaid (red “x” indicates soma position). **c**, Dendritic RF area as a function of dendritic distance to soma; data pooled across cells of the same type (see below). **d**, Comparison of RF area change with dendritic distance to soma between each pair of RGC type. Red shaded areas indicate dendritic sections with significant differences between types (Methods). **e**, Cells from (a) with arrows indicating spatial offset between ROI centre and the RF contour’s geometrical centre, with arrows pointing at the latter. **f**, RF offset of all recorded ROIs as a function of dendritic distance to soma. **g**, Like (d) but for RF offset changes. Data from tOff alpha (n=17\1,452\850 cells\total ROIs\ROIs passing the quality test), tOff mini (n=5\387\295), sOff (n=4\208\154) and F-mini^Off^ RGCs (n=5\265\126); for individual cell morphologies, see Suppl. Fig. S2. Scale bar: a, 50 µm. For statistical analysis, see Suppl. Information.

Synchronization of dendrites can originate from strong backpropagation of somatic spikes to the dendrites (reviewed in (Stuart and Spruston, 2015)). This is not only expected to increase dendritic RF size but should also shift the RF’s centre closer towards the soma. In contrast, for a more isolated dendrite without backpropagation, the RF centre should roughly correspond to the respective ROI position. Therefore, we next measured ROI-to-RF-centre offsets for the four RGC types (Fig. 3e, f). We found that tOff alpha cells displayed small offsets that did not change much as a function of dendritic distance from the soma. In contrast, the other three RGC types displayed large offsets, with the RF centre strongly shifted towards the centre of the dendritic arbour, which in tOff mini and sOff cells also coincided with the soma (Fig. 3e). Moreover, in tOff mini and sOff cells, offsets increased with dendritic distance from the soma (Fig. 3f). In F-mini^Off^ cells, due to their asymmetrical dendritic arbours, offsets increased with dendritic distance from the arbour centre (Fig. 3f), resulting in an inverted-bell shaped curve. For large dendritic distances, the offsets were significantly different between all pairs of RGC types (Fig. 3g). These results confirm that dendrites of tOff mini, sOff and F-mini^Off^ cells are more synchronized than those of tOff alpha cells, possibly due to backpropagation.

Strongly isolated dendrites, as observed in tOff alpha cells, could allow dendritic computations at a finer spatial scale than the whole cell’s RF. Such isolated dendrites are expected to be spatially more independent than the better synchronized dendrites of tOff mini, sOff and F-mini^Off^ cells. To test this prediction, we determined the overlap of RFs for every ROI pair recorded in a single cell (Fig. 4a, b). We then assessed how the overlap changed with dendritic distance and angle between ROIs (Fig. 4b, c; Supplementary Fig. S3). We found localized and spatially independent RFs only in tOff alpha RGCs (Fig. 4a, c). Here, RF overlap decreased substantially with increasing dendritic and angular distance between ROIs, in line with our previous results. In tOff mini cells, RFs showed partial overlap even when the ROIs were located at opposite sides of the dendritic arbour (Fig. 4a, c). For sOff and F-mini^Off^ cells, RFs overlapped substantially, independent of dendritic and angular distance between ROIs. As a result, the RF overlap maps significantly differed between tOff alpha and the other three RGC types, and partially between tOff mini and the remaining two RGC types (Fig. 4d), supporting significant differences in dendritic processing – from more local in tOff alpha to more global in F-mini^Off^.

**Figure 4.**
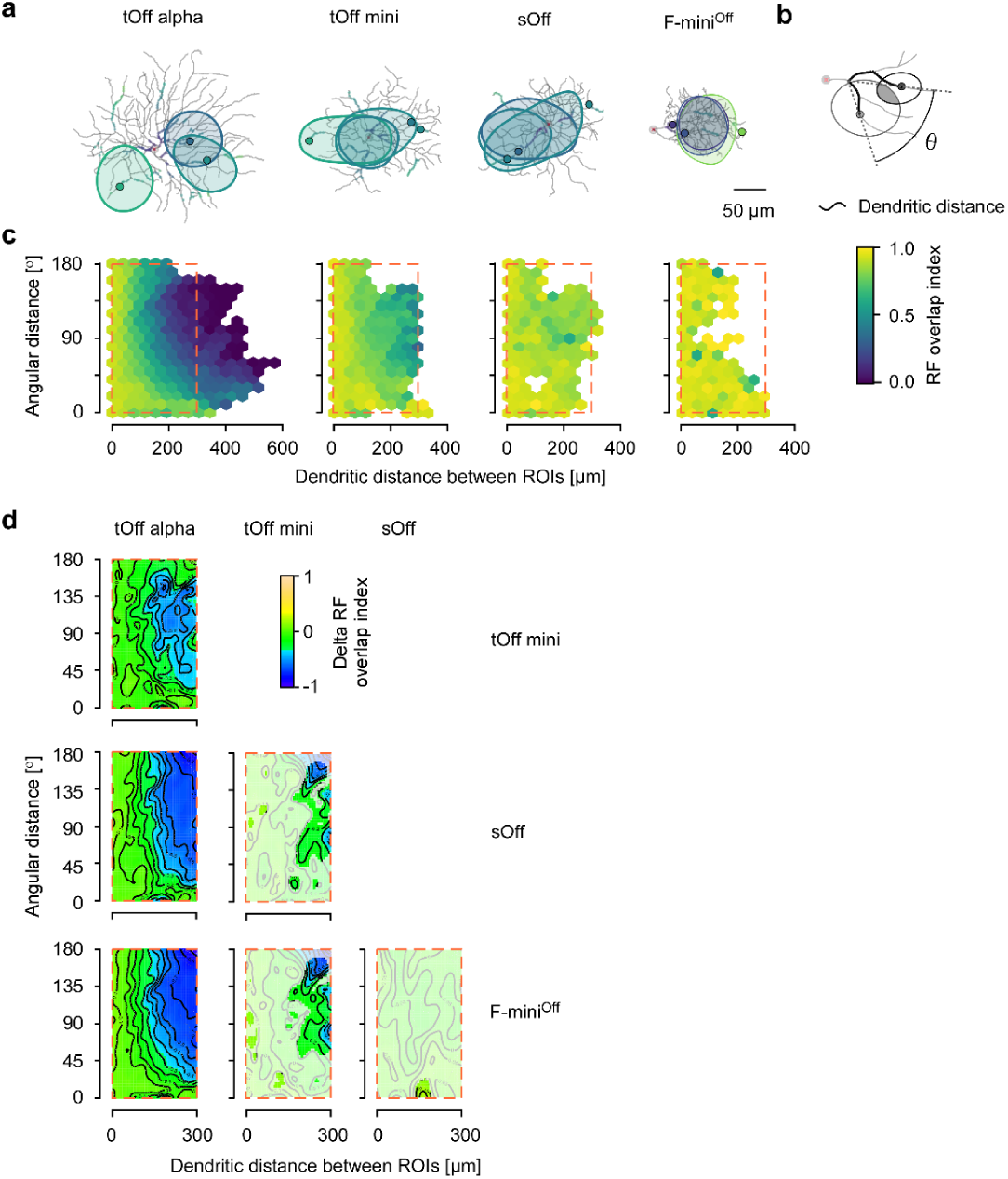
Dendritic RF overlap. **a**, Top-views of the different reconstructed RGC types with RF contours of three ROIs overlaid. **b**, Illustration of dendritic and angular distance (Θ) between two ROIs (measured from the last common branching node) and RF overlap (grey area) of two RF contours (ellipses). **c**, Hexagon maps showing the dendritic RF overlap index (colour-coded) as a function of Θ and dendritic distance for all ROI pairs: tOff alpha (n=17\40,777 cells\ROI pairs), tOff mini (n=5\13,524), sOff (n=4\3,141), and F-mini^Off^ (n=5\2,097). **d**, 2D comparison maps for plot area marked by the dashed red rectangles in (c) for each pair of RGC types. Colour codes difference in RF overlap index, with whitened areas indicating no significant difference. For statistical analysis, see Suppl. Information.

Together, these results suggest that different RGC types with similar input profiles apply vastly different dendritic integration rules. For example, the dendrites of tOff alpha cells seem to exhibit little backpropagation but reasonably strong forward propagation, integrating RFs from all dendrites symmetrically. This leads to larger proximal than distal RFs and distal RFs with little overlap and displacement. In contrast, the other three RGC types show strong indication for backpropagation across their dendritic arbour, causing distal RFs to be highly overlapping and displaced towards the centre of the dendritic arbour.

### Temporal dendritic integration varies between RGC types

Dendritic inputs are not only integrated across space, but also over time. To relate spatial to temporal dendritic integration, we next probed the temporal synchronisation of light responses across the dendritic arbour of the four RGC types. For that, we used a “chirp” stimulus that consisted of a light step followed by frequency and contrast modulations (Methods) and was presented as local (100 µm in diameters) and full-field (500 µm) version. Notably, F-mini^Off^ RGCs did not show any reliable dendritic chirp responses, despite the same ROIs passing our RF quality threshold (Methods). This finding is consistent with earlier observations in this RGC type (*cf*. “x2 cell” of Extended Data Fig. 5 in (Baden et al., 2016)). Therefore, we focussed the following analysis on the remaining three RGC types.

We found that dendritic responses to the local chirp in tOff alpha and tOff mini RGCs were quite similar but differed from those in sOff RGCs (Fig. 5a-c). In the latter, local chirp responses were more sustained than those in the other two types (Supplemental Fig. S4c); this difference resonates with sOff cells stratifying slightly more distally (*cf*. Fig. 2a) and, hence, presumably receiving more input from sustained BC types (Franke et al., 2017; Roska and Werblin, 2001; Yu et al., 2018). When presented with the full-field chirp, tOff alpha and tOff mini RGC responses became somewhat more distinct (i.e. to the frequency modulation). This difference was not found in an earlier study (Baden et al., 2016) but may be related to the fact that in the present study, light stimuli could be precisely centred on the recorded cell. In addition, all three RGC types often showed On events that were much less frequent for the local chirp (Fig. 5a-c; Supplemental Fig. S4b, d). Similar On-events in Off-cells have also been observed in BC responses (Franke et al., 2017). In general, differences between full-field and local chirp responses were more pronounced in sOff RGCs (Supplemental Fig. S4e), suggesting that stimulus size had a larger effect on sOff cells compared to the other two types. This could be due to a stronger inhibitory surround or connections to BCs that are more strongly influenced by surround stimulation (Franke et al., 2017).

**Figure 5.**
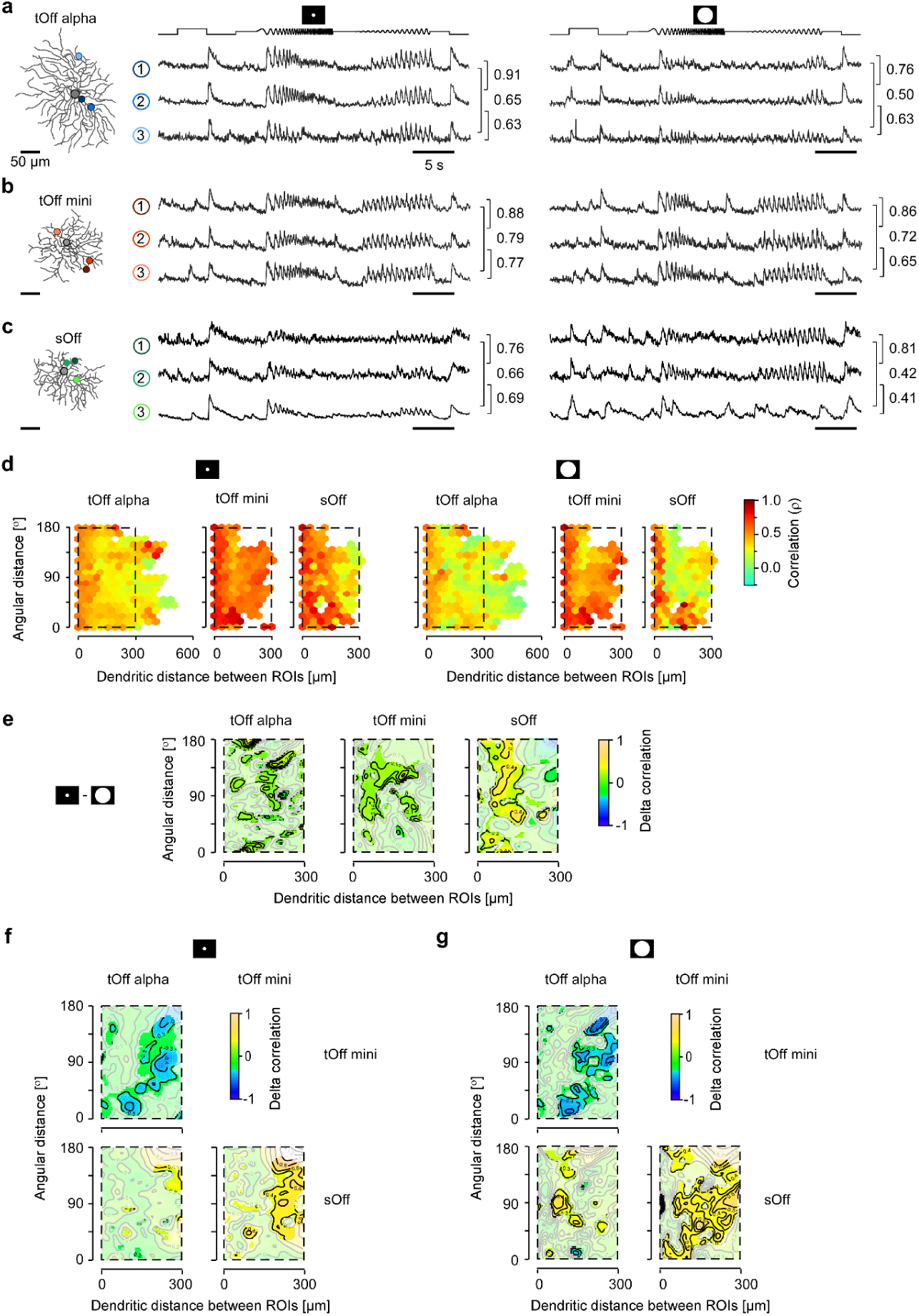
Temporal correlation across dendrites. **a**, Exemplary response of a tOff alpha RGC to local (middle) and full-field chirp (right) recorded from three ROIs indicated on the reconstructed cell (left). Values next to the traces indicate linear correlation coefficient of the corresponding trace pair. **b**,**c**, Like (a), but for tOff mini (b) and sOff RGC (c). **d**, Hexagon maps showing response correlations for local (left) and full-field chirp (right) as a function of angular distance and dendritic distance between ROIs for tOff alpha (n=17\12,770\13,001 cells\pairs for full-field\pairs for local), tOff mini (n=5\6,529\6,529) and sOff RGCs (n=4\2,622\2,557). Colour encodes correlation. **e**, 2D comparison maps for inter-ROI correlation of local and full-field chirp responses for the plot area marked by dashed black rectangle in (d) for each RGC type. Colour codes difference in correlation, with whitened areas indicating no significant difference. **f**,**g**, Like (e), but for the comparison between cell types for local chirp responses (f) and full-field chirp responses (g). For statistical analysis, see Suppl. Information.

To analyse the temporal properties of dendritic integration in these cells, we quantified the correlation of local or full-field chirp responses between ROI pairs across the dendritic arbour (Fig. 5d). In all three RGC types, correlations between ROIs were higher for responses to local than to full-field chirps (Fig. 5d, e). The decorrelation observed for full-field chirps was especially pronounced in sOff cells (Fig. 5e). In tOff alpha and sOff RGCs, correlation decreased with dendritic and angular distance (Fig. 5d). While in tOff alpha cells, lower correlation coincided with smaller and less overlapping RFs, sOff RGCs displayed low correlation in their distal dendrites while at the same time featuring highly overlapping RFs across the whole dendritic arbour. In contrast to the other two RGCs, temporal correlation in tOff mini cells was largely independent of dendritic and angular distance (Fig. 5d). In addition, correlation was overall much higher, indicating that dendritic segments in tOff mini cells are temporally more synchronized (*cf.* Fig. 5c). Because differences in temporal correlation between RGC types persisted when applying a more stringent quality criterion (Supplemental Fig. S6; Methods), it is unlikely that they were due to systematic differences in recording quality (i.e. signal-to-noise-ratio).

Taken together, our data suggest that spatio-temporal integration is tuned across the RGC dendritic arbour in a highly type-specific manner (Fig. 5f, g). The studied RGC types ranged between two main dendritic integration profiles: The first profile featured strongly isolated dendrites (e.g. in tOff alpha) and may render the cell sensitive to fine visual stimulus structures within the cell’s RF. In contrast, the second profile featured strongly synchronised dendrites with highly overlapping RFs (e.g. in tOff mini RGCs) and may tune the cell towards robustly detecting a stimulus independent of its location within the RF.

### Simulation reveals mechanistic principles for type-specific dendritic integration

The dendritic integration properties of RGC types may be influenced by morphological features, such as branching pattern, dendritic thickness and segment length, and the complement and distribution of ion channels (Koch et al., 1982; Rall and Rinzel, 1973). To understand which of these properties may explain the dendritic integration profiles we observed, we built a simple, morphology-inspired biophysical model and focussed on the effects of the type-specific morphology and dendritic channel densities in tOff alpha and tOff mini cells (Fig. 6a).

**Figure 6.**
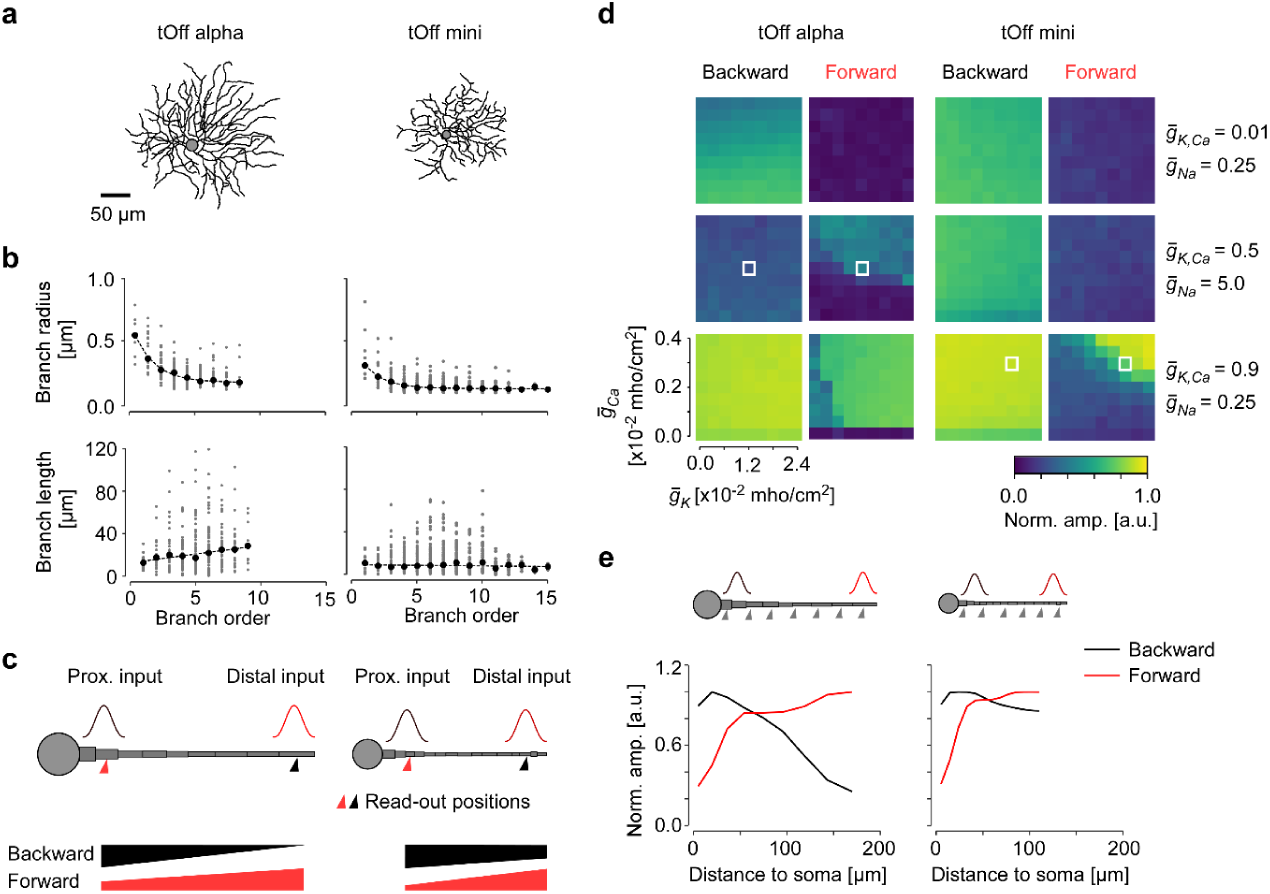
Simulation of dendritic signal propagation in tOff alpha and tOff mini RGCs. **a**, Reconstructed cell morphologies of tOff alpha and tOff mini RGC (same cells as in Figs. 3,4). **b**, Dendrite radius (top) and segment length as functions of branch order (data from http://museum.eyewire.org; n=2 for tOff alpha (4ow); n=3 for tOff mini (4i)). **c**, Illustration of the ball-and-stick models used for simulations in (d, e). Simulated inputs at proximal (25 µm to soma) and distal (85% of the total dendrite length to soma) positions indicated as red and black Gaussians, respectively. Respective read-out positions for (d) are indicated below the dendrite. The thickness change of the bars (bottom) corresponds to the decay of forward (red) and backward (black) signal propagation expected from our experimental data. **d**, Heat maps showing the signal amplitude at the two read-out positions indicated in (c), normalized to the amplitude at the respective input position as a function of ion channel density combinations. White boxes indicate channel combinations that are consistent with our experimental results. **e**, Normalized signal amplitude at read-out positions along the dendrite as a function of dendritic distance for the channel combinations indicated by boxes in (d). Generic voltage-gated 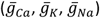 and Ca^2+^ activated 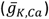 conductances were modeled after Fohlmeister and Miller (1997). For details, see methods.

To capture the morphological differences between the cell types, we first extracted morphological parameters from a published EM dataset (Bae et al., 2018). We found that for tOff alpha cells, dendritic radius decreased systematically with increasing branch order; this decrease was less pronounced in tOff mini cells (Fig. 6b). In addition, dendritic segment length increased with branch order for tOff alpha cells, while it remained constant for tOff mini cells. Based on these differences, we built a ball-and-stick model for each cell type (Fig. 6c). For our simulations, we provided either a proximal or distal input, with readout positions at the dendritic tip and close to the soma (Methods). Based on the dendritic integration profiles of tOff alpha and tOff mini cells (*cf*. Figs. 3-5), we hypothesize that *(i)* in tOff alpha cells, forward propagation (from distal to proximal dendrites) should be stronger than backward propagation and *(ii)* that backpropagation should be strong in tOff mini cells (Fig. 6c).

To investigate the role of ion channel distribution on dendritic signal propagation, we systematically varied the dendritic density of Ca^2+^-activated K^+^ channels 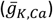 and voltage-gated K^+^, Na^+^ and Ca^2+^ channels (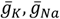 and 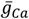). Notably, the same combination of channel densities had quite different effects when applied to the two RGC morphologies (compare columns in Fig. 6d and Supplemental Fig. 7), highlighting how strongly the interplay between morphology and channel complement affects a cell’s dendritic signal propagation. Notably, we found that distinct, cell-type specific sets of ion channel densities were compatible with the experimentally derived hypotheses (Fig. 6d): For the tOff alpha cell model, intermediate 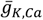 and high 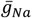 and 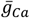 channel densities were required to generate stronger forward propagation compared to backward propagation (Fig. 6d, e). For the same channel densities, forward propagation in modelled tOff mini cell was so low that distal inputs were almost completely extinguished before reaching the proximal dendrite. In contrast, with higher 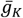 and lower 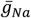 densities, tOff mini cells showed strong backward and substantial forward propagation, in line with our hypothesis (Fig. 6d, e).

Together, these results suggest that morphology alone does not explain the experimentally observed differences between the two cell types. Instead, our model indicates that differences in dendritic channel densities may be responsible for the distinct dendritic integration profiles in RGCs.

## DISCUSSION

Here, we studied dendritic integration in four types of mouse Off RGC (tOff alpha, tOff mini, sOff, and F-mini^Off^), which have their dendrites in overlapping strata of the IPL and, hence, receive highly overlapping sets of synaptic input. Recordings of local, light-evoked dendritic activity and compartmental modelling revealed surprising differences between the cells’ spatio-temporal dendritic integration. What could these distinct integration rules be good for in terms of visual computations?

In tOff alpha RGCs (Deny et al., 2017; Krieger et al., 2017; Pang et al., 2003; Van Wyk et al., 2009), as the distance from the soma increased, RF area decreased and dendritic RFs became increasingly non-overlapping, with minimal offset between recording site and respective RF centre. In addition, activity on different dendritic branches was only moderately correlated. The more isolated, independent dendritic segments in tOff alpha cells may help them to detect fine structures of visual stimuli and support visual computations relying on spatial resolution below the RF of the entire cell. This is reminiscent of what has been reported about On alpha cells, which possess nonlinear RFs and respond to patterns that contain local structures finer than the cell’s RF centre (Schwartz et al., 2012). In contrast, in tOff mini and sOff RGCs (Baden et al., 2016), RFs overlapped extensively and changed little in area, while their centres were systematically shifted towards the soma. In addition, the timing of responses was highly correlated across tOff mini dendrites, suggesting they may reliably detect stimuli independent of their location within the RF. For sOff RGCs, the temporal correlation between the activity of different dendritic branches decreased strongly for larger stimuli, suggesting that the cell’s computational properties change for extended visual stimuli, likely due to lateral amacrine cell circuits kicking in. F-mini^Off^ cells (Rousso et al., 2016) were similar to tOff mini and sOff RGCs with some particularities related to the high asymmetry of their dendritic arbour. Our morphologically inspired biophysical model revealed that morphological difference alone cannot explain these experimentally observed dendritic integration profiles; instead, distinct combinations of morphology, ion channel complements and densities are required.

Dendritic integration rules have been studied extensively in the cortex (e.g. (Schmidt-Hieber et al., 2017; Tran-Van-Minh et al., 2016; Vetter et al., 2001)). In the retina, mainly interneurons have been at the centre of interest: For example, it has been suggested that horizontal cells (Chapot et al., 2017) and A17 amacrine cells (Grimes et al., 2010) provide locally computed feedback by confining signals within single varicosities. Likewise, starburst amacrine cell dendrites compute the direction of motion “dendrite-wise” by dividing their dendritic arbour into isolated sectors which contain 15-20 varicosities each (Euler et al., 2002; Koren et al., 2017; Poleg-Polsky et al., 2018). In RGCs, dendritic integration has been studied in direction-selective (DS) RGCs, where intrinsic properties of their dendritic arbour (Schachter et al., 2010; Sivyer and Williams, 2013) and partially their asymmetry (Trenholm et al., 2011) contribute to the generation of DS output. Reminiscent of our findings in tOff alpha cell, the dendritic arbour of DS RGCs is functionally partitioned, with the “DS mechanism” replicated across the dendritic arbour, such that local motion within the cell’s RF can cause a robust spiking response (Barlow and Levick, 1965; Oesch et al., 2005).

We chose to focus on four types of Off RGCs because they are expected to receive similar excitatory synaptic inputs. Nevertheless, due to small differences in dendritic stratification depth, they make connections with partially different sets of BCs: tOff alpha cells contact mainly transient type 3a and 4 BCs, while sOff cells likely contact mainly the more sustained type 1 and 2 BCs (Bae et al., 2018; Helmstaedter et al., 2013; Yu et al., 2018). In line with this, we found that the dendrites of tOff alpha cells exhibited more transient responses than those from sOff cells. Since tOff mini RGCs co-stratify with tOff alpha RGCs, they likely receive excitatory inputs from the same BC types and thus should exhibit similar response properties. Indeed, these two RGC types showed similar responses to local chirps. Nevertheless, they may be differentially modulated by type-specific connectivity to ACs. In line with this, the two cell types showed more distinct responses to full-field chirps. In principle, the interaction of excitation from BCs and inhibition from ACs may attenuate the excitatory inputs and affect dendritic integration (Roska et al., 2006), raising the possibility that the observed type-specific differences could at least partially result from type-specific microcircuit connectivity rather than mainly from cell intrinsic properties as suggested above. While we cannot exclude this possibility, our simulation results suggest a more parsimonious explanation for the observed differences in these dendritic integration profiles, relying on cell-intrinsic mechanisms only.

Apart from contributions of the microcircuit connectivity, dendritic integration is mainly determined by a combination of morphological features and passive and active membrane properties, which can differ significantly between RGC types (reviewed in (Stuart and Spruston, 2015)). In some RGC types like the tOff alpha, for instance, the dendritic diameter becomes smaller and dendritic segment length gets longer with increasing branch order. This, in turn, results in a higher axial resistance and shorter propagation distance for more distal dendritic signals. In other RGC types like tOff mini, however, dendritic diameter and segment length does not systematically change with increasing branch order. In addition, a variety of ion channels, including Ca^2+^-activated K^+^ channels, hyperpolarization-activated cyclic nucleotide-gated (HCN) channels, and voltage-gated K^+^, Na^+^ and Ca^2+^ channels, have been found in RGC dendrites, differing in density and dendritic locations between cell types (Van Hook et al., 2019).

An earlier theoretical study suggested that alpha RGCs – with their large dendritic arbours, thick and short proximal but thin and long distal branches (Wässle et al., 1981) – feature independent dendritic regions (Koch et al., 1982). In contrast, RGCs with constant dendritic diameter and branch length across their dendritic arbour are thought to produce densely coupled dendritic regions. In these RGCs, their morphology could enable more efficient dendritic backpropagation and therefore lead to the synchronization of dendritic signals (Tran-Van-Minh et al., 2015). Indeed, we observed more independent dendritic regions in tOff alpha cells, but more spatially synchronized dendritic regions in tOff mini, sOff and F-mini^Off^ cells. In tOff mini and tOff alpha cells, their forward and backward propagation were differentially modulated by the same combinations of ion channel densities, confirming that dendritic morphology is a key determinant of dendritic signal propagation efficiency. However, our simulation results suggest that the dendritic integration properties of tOff alpha and tOff mini RGCs could not be explained by dendritic morphology alone but require dendritic ion channels in agreement with earlier simulation studies (Maturana et al., 2014). One possible reason might be that for most RGCs, action potentials generated in the soma can back propagate to the dendritic arbour (Velte and Masland, 1999), which needs dendritic ion channels to enable the efficient backpropagation (van Rossum et al., 2003; Velte and Masland, 1999).

Our simulation results are based on highly simplified ball-and-stick models of RGC dendrites, as these allowed us to focus on the principles of dendritic integration. Obviously, these models come with several caveats and possibilities for future extensions: First, the morphological parameters we used were extracted from EM data (Bae et al., 2018), where tissue can shrink due to chemical fixation, such that we may have underestimated the axial conductance based on dendrite diameter. Second, it has been reported that the branching pattern is an important variable for determining propagation efficiency of dendritic signals, mainly because of the diameter changes at branch-points (Ferrante et al., 2013). Third, the density and complement of ion channels can vary along the dendrite (Hausser et al., 2000; Van Hook et al., 2019), raising the possibility that spatially varying ion channel densities would allow for more refined control over dendritic computations. Finally, dendritic signalling is driven by the complex interaction of excitatory and inhibitory inputs (as already mentioned above) and the locations of the respective synapses, which will require more precise connectomic studies of the cell types and microcircuits in question. A more realistic model incorporating these aspects could allow additional insights into the mechanisms underlying the observed spatio-temporal dendritic integration rules.

## METHODS

### Animals and tissue preparation

Mice used in this study were purchased from Jackson Laboratory and housed under a standard 12 hour day/night cycle. For all experiments, mice aged 5-8 weeks of either sex were used. We used the transgenic mouse line B6;129P2-*Pvalb*^*tm1(cre)Arbr*^/J (“PV”, JAX 008069, The Jackson Laboratory, Bar Habor, ME; (Hippenmeyer et al., 2005)) cross-bred with the red florescence Cre-dependent reporter line Gt(ROSA)26Sor^tm9(CAG-tdTomato)Hze^ (“Ai9^tdTomato^”, JAX 007905) for all recordings of tOff mini, sOff and F-mini^Off^ cells (n=25 animals). For alpha RGC recordings, we also used the wild-type line (C57Bl/6J, JAX 000664, n=3 animals), as alpha RGCs can be easily targeted due to their large soma size. All animal procedures followed the laws governing animal experimentation issued by the German federal government and were approved by the institutional animal welfare committee of the University of Tübingen.

Mice were dark adapted ≥ 2 hours before tissue preparation, then anaesthetized with isoflurane (Baxter, Hechingen Germany) and killed with cervical dislocation. The eyes were quickly enucleated in carboxygenated (95% O2, 5% CO2) artificial cerebral spinal fluid (ACSF) solution containing (in mM): 125 NaCl, 2.5 KCl, 2 CaCl2, 1 MgCl2, 1.25 NaH2PO4, 26 NaHCO3, 20 glucose, and 0.5 L-glutamine (pH 7.4). After removing cornea, sclera and vitreous body, the retina was flattened on an Anodisc (0.2 µm pore size, GE Healthcare, Pittsburgh, PA) with the ganglion cell side facing up and then transferred to the recording chamber of the microscope, where it was continuously perfused with carboxygenated ACSF (at 35°C and 4 ml/min). All experimental procedures were carried out under very dim red light.

### Loading of individual cells with calcium indicator

To visualize blood vessels and avoiding them when filling individual RGCs, 5 µl of a 50 mM sulforho-damine-101 (SR101, Invitrogen /Thermo Fisher Scientific, Dreieich, Germany) stock solution was added per litre ACSF solution. Sharp electrodes for single-cell injection were pulled on a P-1000 micropipette puller (Sutter Instruments, Novato, CA) with resistances ranging between 70 and 130 MΩ. Oregon Green BAPTA-1 (OGB-1, hexapotassium salt; Life Technologies, Darmstadt, Germany; 15 mM in water) was loaded into individual RGCs using the single-pulse function (500 ms, −10 nA) of a MultiClamp 900A amplifier (Axon Instruments/Molecular Devices, Wokingham, UK). To allow the cells to completely fill and recover, we started recordings 1 h post-injection.

### Two-photon imaging and light stimulation

A MOM-type two-photon microscope (designed by W. Denk, MPI, Martinsried; purchased from Sutter Instruments/Science Products) as described previously (Euler et al., 2009) was used for this study. Briefly, the system was equipped with a mode-locked Ti:Sapphire laser (MaiTai-HP DeepSee, Newport Spectra-Physics, Darmstadt, Germany), green and red fluorescence detection channels for OGB-1 (HQ 510/84, AHF, Tübingen, Germany) and SR101/tdTomato (HQ 630/60, AHF), and a water immersion objective (W Plan-Apochromat 20x/1,0 DIC M27, Zeiss, Oberkochen, Germany). For all scans, we tuned the laser to 927 nm, and used a custom-made software (ScanM, by M. Müller, MPI, Martinsried, and T.E.) running under IGOR Pro 6.3 for Windows (Wavemetrics, Portland, OR). Time-elapsed dendritic signals were recorded with 64×16 pixel image sequences (31.25 Hz). High-resolution morphology stacks were acquired using 512×512 pixel image stacks with 0.8 or 1.0 µm z steps.

Light stimuli were projected through the objective lens (Euler et al., 2009). We used two alternative digital light processing (DLP) projectors: a K11 (Acer, Ahrensburg, Germany) or a LightCrafter E4500 MKII (Texas Instruments, Dallas, TX; modified by EKB Technologies Ltd., Israel). Both were equipped with light-emitting diodes (LEDs) – “green” (575 nm) and UV (390 nm) that match the spectral sensitivities of mouse M- and S-opsins (for details, see (Baden et al., 2016; Franke et al., 2019)). Both LEDs were intensity-calibrated to range from 0.1 ×10^3^ (“black” background) to 20.0 ×10^3^ (“white” full field) photoisomerisations P*/s/cone. The light stimulus was centred before every experiment, ensuring that its centre corresponded to the centre of the microscope’s scan field. For all experiments, the tissue was kept at a constant mean stimulator intensity level for ≥ 15 s after the laser scanning started and before light stimuli were presented.

Light stimuli were generated and presented using the Python-based software package QDSpy (see Table 4). Three types of light stimuli were used:

**Table 2.** Model parameters.

| Parameters | Values |
| --- | --- |
| Temperature | $T = 32^\circ\text{C}$ |
| Intracellular axial resistivity | $R_a = 110 \Omega \cdot \text{cm}$ |
| Specific membrane resistance | $R_m = 15000 \Omega \cdot \text{cm}^2$ |
| Specific membrane Capacitance | $C_m = 1 \mu\text{F}/\text{cm}^2$ |
| Potassium reversal potential | $V_K = -75 \text{ mV}$ |
| Sodium reversal potential | $V_{Na} = 35 \text{ mV}$ |

**Table 3.** Reference distribution of ion channels in cell compartments.

| Channel type | Conductance<br>in [S/cm<sup>2</sup>] |  |
| --- | --- | --- |
|  | Soma | Dendrites |
| $\bar{g}_{Na}$ | 0.08 | 0.025 |
| $\bar{g}_{Ca}$ | 0.0015 | 0.002 |
| $\bar{g}_K$ | 0.018 | 0.012 |
| $\bar{g}_{K,A}$ | 0.054 | 0.036 |
| $\bar{g}_{K,Ca}$ | 0.000065 | 0.000001 |

**Table 4.** Software used and repositories for custom scripts and data.

| Part | Description (link) | Company /Author | Item number (RRID) |
| --- | --- | --- | --- |
| ScanM | 2P imaging software running under IGOR Pro | Written by M. Müller (MPI Neurobiology, Martinsried), and T.E. |  |
| IGOR Pro (v6) | https://www.wavemetrics.com | Wavemetrics, Lake Oswego, OR | IGOR Pro v6 (SCR_000325) |
| Python (v3.6.7) | http://www.python.org/ |  |  |
| R (v2.3) | The R project<br>http://www.r-project.org/ |  |  |
| QDSpy (v0.77) | Visual stimulation software<br>https://github.com/eulerlab/QDSpy | Written by T.E, supported by Tom Boissonnet (EMBL, Monterotondo) | - (SCR_016985) |
| Scikit-learn (v0.20.0) | Software package for Python | (Pedregosa et al., 2011) |  |
| Scikit-Image (v0.14.2) | Software package for Python | (van der Walt et al., 2014) |  |
| itsadug (v2.3) | Software package for R | (van Rij et al., 2017) |  |
| mgcv (v1.8-24) | Software package for R | (Wood, 2006) |  |
| NEURON (v7.7.0) | Simulation environment for modelling individual neurons and networks of neurons | (Carnevale and Hines, 2006) |  |
| Custom scripts and data | http://retinal-functomics.net/ |  |  |

1. binary dense noise (20×15 matrix of 30 μm/pixel; each pixel displayed an independent, balanced random sequence at 5 Hz for 5 minutes) for spatio-temporal receptive field (RF) mapping;
2. full-field (800×600 µm) “chirp”, consisting of a bright step and two sinusoidal intensity modulations, one with increasing frequency (0.5-8 Hz) and one with increasing contrast.
3. local “chirp”; like (2) but with a diameter of 100 µm.

### Reconstruction of cell morphologies

Directly after the recording, the complete dendritic morphology of the RGC was captured by acquiring a high-resolution stack. In case the cell was not bright enough to see all branches in detail, a second dye-injection was performed. Using semi-automatic neurite tracing (Longair et al., 2011), we obtained cell skeletons of the recorded RGCs. If necessary, we de-warped image stack and traced cell, as described earlier (Baden et al., 2016). All further analysis, such as the extraction of morphological parameters (see below), was done using custom Python scripts.

### Relating recording positions to cell morphology

As the full dendritic morphology could not be imaged during calcium recordings, recorded dendrites (i.e. ROIs, see below) were not necessarily well-aligned with the cell morphology reconstructed later. Based on the relative position of each recording field, a region 9 times larger than this recording field was cropped from the reconstructed morphology and z-projected. Next, the recording field was automatically aligned to this cropped region using `match_templates` from *scikit-image* (Table 4). The centre coordinates of all ROIs in that recording field were then calibrated to the closest dendritic branch based on their Euclidean distance. In rare cases, when automatic matching failed, the matching was done manually.

### Morphological parameters and hierarchical clustering

To morphologically cluster the RGCs as described by Bae et al. (2018), we had to determine the relative position of the two ChAT bands, the dendritic plexi of the starburst amacrine cells (Vaney, 1984). For this, the blood vessel plexi in GCL and INL served as landmarks. With their positions defined as 0 and 1, the relative IPL depth of the On and Off ChAT bands is 0.48 and 0.77, respectively, as shown earlier (Baden et al., 2016; Franke et al., 2017). The following parameters were extracted for each cell:

To determine the marginal-central *arbour density difference*, we defined the central IPL as the portion between the ChAT bands and the remainder (On ChAT band to GCL, Off ChAT band to INL) as marginal IPL (*cf*. Fig. 2b, inset). The marginal-central arbour density difference was calculated using the sum of the dendritic length in central IPL minus the sum of the dendritic length located in marginal IPL.

*Dendritic arbour area* was calculated as the area of the tightest convex hull containing the z-projected dendritic arbour.

*Asymmetry* of the dendritic arbour was calculated as the distance between the centre of mass of dendritic density and the soma position.

*Soma size* was defined as soma area. For this, the image frame in which the soma appeared the largest was used.

*Dendritic distance between ROIs* was defined as the shortest distance along the dendrite between two ROIs.

*Angular distance between ROIs* was defined as the positive angle between two ROIs and the nearest branching point (*cf*. Fig. 4b).

Hierarchical clustering was performed with 1D *k*-means clustering with *k*=2 for all splits, using `KMeans` from the Python package *scikit-learn* (Table 4). First, cells were split into two clusters based on arbour density difference (*cf*. Fig. 2b). Next, the group with lower arbour density difference was separated by soma size, while the group with the higher arbour density difference was further split based on their asymmetry index. Here, we refrained from further splitting, because the cells in each group displayed highly consistent light responses. Thus, these four groups were used for further analysis.

### Data analysis

#### All data were analysed using custom scripts

For data preprocessing, we used IGOR Pro; further analysis and modelling was done using Python and R. Upon publication, all data, scripts and models will become available (see links in Table 4).

#### Regions of interest (ROIs)

We used dense noise recordings to extract ROIs. First, for each recorded field, the standard deviation (s.d.) of the fluorescence intensity for each pixel over time was calculated, generating an s.d. image of the time-lapsed image stack. Pixels brighter than the mean of the s.d. image plus 1 s.d. were considered dendritic pixels. Then, in each recorded field, the time traces of the 100 most responsive dendritic pixels (=100 brightest dendritic pixels in the s.d. image) were extracted and cross-correlated. The mean of the resulting cross-correlation coefficients (*ρ*) served as correlation threshold (*ρ*_*Threshold*_) for each field. Next, we grouped neighbouring pixels (within a distance of 3 µm) with *ρ* > *ρ*_*Threshold*_ into one ROI. Finally, each ROI’s Ca^2+^ trace was extracted using the image analysis toolbox SARFIA for IGOR Pro (Dorostkar et al., 2010). A time marker embedded in the recorded data served to align the traces relative to the visual stimulus with 2 ms precision. All stimulus-aligned traces together with the relative ROI positions on the recorded cell’s dendritic arbour were exported for further analysis.

#### Dendritic receptive fields

Dendritic RFs were estimated using Automatic Smoothness Determination (ASD, (Sahani and Linden, 2003)), a linear-Gaussian encoding model within the empirical Bayes framework. The relationship between stimulus and response was modelled as a linear function plus Gaussian noise:

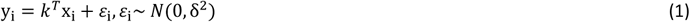

where x is the binary dense noise stimulus (20×15 matrix of 30 µm/pixel), y is the gradient of the Ca^2+^ response, *k* is the spatio-temporal RF (STRF) with a time lag ranging from −1,000 ms to 0 ms, and *ε* is independent and identically distributed (i.i.d.) Gaussian noise with zero mean and δ^2^ variance.

The STRF was then calculated in two steps (Sahani and Linden, 2003): First, the ASD prior covariance (*C*_*ij*_ = exp(-ρ-Δ_*ij*_/2δ), where Δ_*ij*_ is the squared distance between any two filter coefficients), controlled by the spatial and temporal smoothness (δ) and scale (ρ), was optimized using evidence optimization. Then, the STRF was estimated by maximum a posteriori linear regression between response and stimulus using the optimized prior. The spatial RF maps shown represent the spatial component of the singular value decomposition of the STRF.

To quantify the quality of spatial RFs, contours were drawn on up-sampled and normalized RF maps with different thresholds (0.60, 0.65 and 0.70). The quality was then determined from the number of contours, their sizes and their degree of irregularity. The irregularity index was defined as

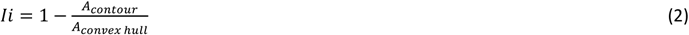

with *A*_*contour*_ corresponding the area of the RF contour, and *A*_*convex hull*_ the 2D morphology’s convex hull. Only data with a good RF (a single contour with *Ii* < 0.1, *A*_*contour*_> 1.8 . 1,000µm^2^, at a contour threshold of 0.60; see Suppl. Fig. S1) were used for further analyses.

#### Offset between ROI centre and its RF centre

RF offset was calculated as the linear distance between ROI centre and the geometrical centre of its RF contour.

#### Dendritic RF overlap index

An RF overlap index (*Oi*) was calculated as follows:

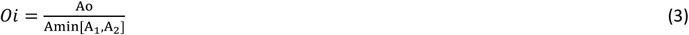

where *A*_1_ and *A*_2_ are the RF areas of the ROI pair, Ao is the overlap area between *A*_1_ and *A*_2_, and *Amin*[*A*_1,_ *A*_2_] corresponds to the smaller area (*A*_1_ or *A*_2_).

#### Full-field chirp and local chirp

Ca^2+^ traces for full-field and local chirp stimuli were linearly up-sampled (interpolated) to 500 Hz, baseline-subtracted (using the mean of 2,500 samples before light stimulus onset) and normalized by the s.d. of this baseline. To estimate the signal-to-noise ratio, we calculated the response quality index (*Qi*) for both full-field and local chirps as described before (Franke et al., 2017):

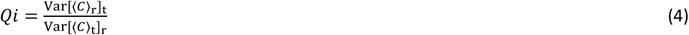

where *C* is the T-by-R response matrix (time samples by stimulus repetitions) and ⟨⟩ _x_ and Var[] _x_ denote the mean and variance across the indicated dimension, respectively. If all trials were identical, such that the mean response is a perfect representative of the response, *Qi* = 1. If all trials were completely random with fixed variance, such that the mean response is not informative about the individual trials, *Qi* ∝ 1/*R*.

#### Signal correlation

To quantify temporal signal correlation, we cross-correlated the mean Ca^2+^ responses. We noticed that some ROIs with good spatial RFs (see above) displayed a low signal-to-noise chirp responses. Hence, we repeated the analysis for *Qi* > 0.4 or 0.5 with comparable result (Supplemental Fig. S6).

#### Further temporal analysis

Using the responses to the step part of the chirp stimulus, we calculated a transience index (*Ti*, for local chirp) and a polarity index (*POi*, for both local and full-field chirps). Here, only ROIs with *Qi* > 0.4 were used for the analysis. Before the computation of these indices, the mean traces were binomially smoothed (with 3,000 repetitions). Then, 2 s.d. of the baseline (2,500 samples of the smoothed trace before light stimulus onset) were used to determine the time of response onset (*T*_*R_onset*_) and offset (*T*_*R_offset*_).

*Ti* was calculated as

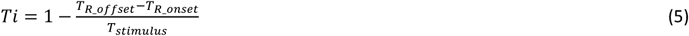

where *T*_*stimulus*_ is the stimulus duration.

For *POi*, data points before and after the response (see above) were set to zero, before calculating:

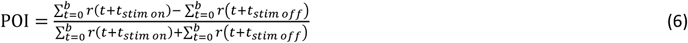

Where *b* = 3 s, *t*_*stim on*_ and *t*_*stim off*_ are the time points of light stimulus onset and offset, *r*(*t*) is the mean response at time *t*. For ROIs responding only to the light-onset, *POi* = 1, whereas for ROIs only responding during the light-offset, *POi* = −1.

### Statistical analysis

We used Generalized Additive Models (GAMs) to analyse the relationships of RF size *vs*. dendritic distance; RF offset vs. dendritic distance; RF overlap vs. dendritic distance and dendritic angle; temporal correlation vs. dendritic distance and dendritic angle (for details, see Supplemental Information). GAMs extend the generalized linear model by allowing the linear predictors to depend on arbitrary smooth functions of the underlying variables (Wood, 2006):

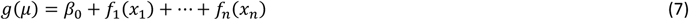

Here, *x*_*i*_ are the predictor variables, *g* is a link function, and the *f*_*i*_ are smooth functions of the predictor variables. These smooth functions can also depend on more than one predictor variable.

To implement GAMs and perform statistical testing, we employed the *mgcv* package for R (Table 4). Here, for smooth terms we used penalized regression splines. We modelled the dependence of our variable of interest as a single smooth term per cell type for univariate dependencies and a tensor product smooth for bivariate dependencies. The dimension of the basis was set high enough such that the estimated degrees of freedom stayed sufficiently below the possible maximum. Smoothing parameters were selected via the default methods of the package. All models also included a random effect term for cell identity. Typically, we used models from the Gaussian family; for the dependence of RF overlap on dendritic distance and dendritic angle, we instead used a scaled t-distribution, as this improved BIC (Bayesian information criterion) and diagnostic plots.

Statistical significance for differences in the obtained smooths between cell types were performed using `plot_diff` or `plot_diff2` of the *itsadug* package for R (Table 4). 95% confidence intervals were calculated using the simultaneous confidence intervals (CI) option, excluding the random effect of cell identity.

### Biophysical model

To explore the mechanisms underlying dendritic integration in different RGC types, we built a multi-compartmental 1D model. To get precise measurements of dendrite thickness and segment length for tOff alpha and tOff mini cells, we extracted these information from published morphologies of “4ow” and “4i” RGCs (*cf*. Fig. 6b), respectively, reconstructed from EM data (http://museum.eyewire.org). Then we mapped the medium values of these parameters to the respective branch order of the model (*cf*. Fig. 6c). The model was implemented in the NEURON simulation environment (Hines and Carnevale, 1997). Here, each dendritic portion (between two branch points) was represented as a “section” in the simulator, which was further divided into multiple “segments” (compartments) with a maximal length of 7 µm. The 1D model can be characterized by the cable equation,

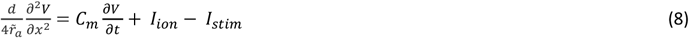

where *V* is the voltage across the cell membrane, *x* is the distance along the cable, *d* is the dendritic diameter, 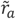 is the intracellular resistivity, and *C*_*m*_ is the specific membrane capacitance. *I*_*ion*_ represents the sum of four voltage-gated cation currents (sodium, *I*_*Na*_; calcium, *I*_*Ca*_ delayed rectifier potassium, *I*_*K*_; A-type potassium, *I*_*K,A*_), one calcium-activated potassium current (*I*_*K,Ca*_), and one leak current (*I*_*Leak*_). The current dynamics are described following Fohlmeister and Miller (1997) as:

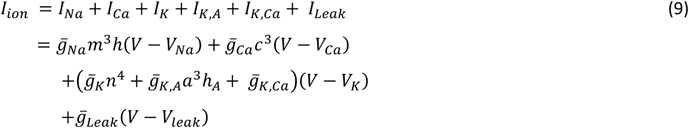

The intracellular stimulation current (*I*_*stim*_ in Eq. 8) was the product of a 5,000 ms x 200 µm 1D Gaussian noise stimulus and a BC’s spatial RF with a Gaussian shape (with the width set by σ=6 and the centre depends on the current injection location). The stimulation current was injected either at the proximal (at 25 µm from soma) or distal dendrite (at 85% of the total dendrite length from soma), then filtered by a soft rectification function. The magnitude of *I*_*stim*_ was scaled between 0 and 15 nA with a mean of 6.17 nA and an s.d. of 1.88 nA to ensure that the input stimulus would elicit spikes in the soma for all parameter combinations.

Model parameters (Table 2) and channel conductances (Table 3) were taken from Fohlmeister and Miller (1997). To identify parameters that explain the experimental data, we grid-searched combinations of ion channel densities by multiplying the reference parameters with different scaling factors (for 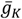 and 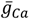, the scaling factors ranged from 0 to 2, and were incremented by 0.25 each step; for 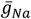, we used [0.1, 1, 2], for 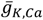, [10, 100, 1000, 5000, 9000]).

Voltage changes in the dendrites were read out at the centre of each dendritic “section” and used to estimate the local dendritic RF with a maximum likelihood method, then smoothed by a Savitzky-Golay filter (window length of 31; 3^rd^ degree polynomial). The peak amplitudes of dendritic RFs were measured and normalized to the RF at the position closest to the current input.

## ACKNOWLEDGEMENTS

We thank Huayu Ding for helping us with single-cell microinjections, Luke Rogerson for help with statistics und discussion, Zhijian Zhao and Gordon Eske for excellent technical support, and Timm Schubert for discussion.

This research was supported by Deutsche Forschungsgemeinschaft (DFG, EXC307 to TE and EXC 2064, project number 390727645 to PB; BE5601/4-1, BE5601/6-1 to PB; EU 42/10-1 to TE); NINDS of the National Institutes of Health (U01NS090562 to TE); BMBF (01GQ1601 and 01IS18052C to PB; 01GQ1002 to KF); BWSF (AZ 1.16101.09 to TB); MPG (M.FE.A.KYBE0004 to KF).

