## Supplementary Information for "Type-specific dendritic integration in mouse retinal ganglion cells"

**Ran, Huang et al.**

- Supplementary Statistical Analysis
- Supplementary Figures
- Supplementary References

### SUPPLEMENTARY STATISTICAL ANALYSIS

**Receptive field size vs. dendritic distance**

To model the dependence of receptive field (RF) size as a function of dendritic distance, we used a Gaussian GAM with factor cell type, a smooth term for each type as a function of dendritic distance and random factor cell id.

```
rf_size ~ type + s(soma_dist, by = type, k = 50) + s(exp_date, bs = "re")
```

We set the basis dimension k=50 to allow for sufficiently “wiggly” smooth terms. Inspection of the model fit indicated that this was high enough (using `gam.check`). The resulting model was fit using n=1,324 data points and yielded the following results:

```
Parametric coefficients:
               Estimate Std. Error t value Pr(>|t|)
(Intercept)     12.7394     0.9679  13.162  < 2e-16 ***
typesustained   -1.4720     2.2130  -0.665  0.50606
typemini alpha  -2.2966     2.0233  -1.135  0.25656
typef-mini      -5.8777     2.0320  -2.893  0.00389 **

Approximate significance of smooth terms:
                                   edf Ref.df      F  p-value
s(soma_dist):alpha transient 10.628 13.241 21.584  < 2e-16 ***
s(soma_dist):sustained        4.821  6.094  2.634 0.014754 *
s(soma_dist):mini alpha       1.599  2.015  7.568 0.000535 ***
s(soma_dist):f-mini           1.806  2.265  0.507 0.564331
```

Thus, the smooth terms for the tOff alpha RGC and the mini alpha RGC are highly significant, indicating non-random variation of receptive field size with dendritic distance.

Overall, the model explained 54.6% of the deviance.

Analysis of the pairwise differences between cell types indicated that the RF size of tOff alpha RGC dendrites close to the soma (< 50 to 75 µm) was significantly larger than that of other RGC types.

**Receptive field offset vs. dendritic distance**

To model the dependence of RF offset as a function of dendritic distance, we used a Gaussian GAM with factor cell type, a smooth term for each type as a function of dendritic distance and random factor cell id.

```
offset ~ type + s(soma_dist, by = type, k = 50) + s(exp_date, bs = "re")
```

We set the basis dimension k=50 to allow for sufficiently “wiggly” smooth terms. Inspection of the model fit indicated that this was high enough (using `gam.check`). The resulting model was fit using n=1,324 data points and yielded the following results:

```
Parametric coefficients:
               Estimate Std. Error t value Pr(>|t|)
(Intercept)      24.396      2.602   9.376  < 2e-16 ***
typesustained    23.990      6.039   3.973 7.51e-05 ***
typemini alpha   12.717      5.415   2.348    0.019 *
typef-mini        4.088      5.493   0.744    0.457
```

```
Approximate significance of smooth terms:
                                     edf Ref.df      F  p-value    
s(soma_dist):typealpha transient  3.341  4.227 11.78 1.15e-09 ***
s(soma_dist):typesustained        9.686 12.062 53.49  < 2e-16 ***
s(soma_dist):typemini alpha       4.349  5.513 50.12  < 2e-16 ***
s(soma_dist):typef-mini           4.114  5.115 23.18  < 2e-16 ***
```

Thus, the smooth terms for all RGC types are highly significant, indicating non-random variation of RF centre offset with dendritic distance.

Overall, the model explained 63.2% of the deviance.

**Receptive Field Overlap Vs. Dendritic Distance and Dendritic Angle**

To model the dependence of RF overlap as a function of dendritic distance and dendritic angle, we used a t-distributed GAM with 5 degrees of freedom. In this case, we found that allowing for t-distributed residuals improved the quality of the model (AIC: -61,825 vs. -59,202).

We included a factor cell type, a bivariate smooth term as a function distance and angle, unique for each type, and a random effect term for cell id.

```
overlap ~ type + te(roi_dist, angle, by=type, k=20) + s(cell_id, bs="re")
```

We set the basis dimension k=20. Inspection of the model fit indicated that this was high enough (using `gam.check`).

The resulting model was fit on n=54,194 data points and yielded the following results:

```
Parametric coefficients:
               Estimate Std. Error t value Pr(>|t|)    
(Intercept)     0.53763    0.02234  24.063  < 2e-16
typesustained   0.28248    0.05067   5.575 2.49e-08
typemini alpha  0.20049    0.04854   4.130 3.63e-05
typef-mini      0.28648    0.04725   6.063 1.34e-09

Approximate significance of smooth terms:
                                     edf Ref.df       F p-value    
te(roi_dist,angle):alpha transient 239.02 291.20 602.962  <2e-16 ***
te(roi_dist,angle):sustained        69.23  95.47   8.983  <2e-16 ***
te(roi_dist,angle):mini alpha      199.26 244.15  46.514  <2e-16 ***
te(roi_dist,angle):f-mini           14.02  19.21   1.884   0.011 *
```

This indicated that all cell types show significant variation in the smooth surface of RF overlap vs. angle and distance.

Overall, the model explained 72.8% of the deviance.

**Correlations vs. dendritic distance and dendritic angle**

To model the dependence of temporal correlation as a function of stimulus (local or global), dendritic distance and dendritic angle, we used a Gaussian GAM with factor cell type, stimulus, and a bivariate smooth term as a function distance and angle, unique for each combination of type and stimulus, and a random effect term for cell id.

```
corr ~ typeXchirp + te(roi_dist, angle, by=typeXchirp, k=50) +
s(cell_id, bs="re")
```

We set the basis dimension k=50. Inspection of the model fit indicated that this was high enough (using `gam.check`).

The resulting model was fit on n= 33,796 data points and yielded the following results:

```
Parametric coefficients:
                                       Estimate Std. Error t value Pr(>|t|)
(Intercept)                            0.418720   0.053359   7.847 4.38e-15 ***
sustained.global                 -0.094504  0.105581  -0.895   0.3707
mini alpha.global                0.213602   0.115928   1.843   0.0654 .
alpha transient.local            0.033782   0.003665   9.217  < 2e-16 ***
sustained.local                  0.028753   0.105457   0.273   0.7851
mini alpha.local                 0.267182   0.115465   2.314   0.0207 *
---

Approximate significance of smooth terms:
                                        edf     Ref.df  F      p-value
te(roi_dist,angle):alpha transient.global 246.85  345.0  11.307  <2e-16 ***
te(roi_dist,angle):sustained.global       190.18  265.0  12.363  <2e-16 ***
te(roi_dist,angle):mini alpha.global      226.69  316.9  10.646  <2e-16 ***
te(roi_dist,angle):alpha transient.local  164.80  234.3  12.321  <2e-16 ***
te(roi_dist,angle):sustained.local        229.01  314.3   9.953  <2e-16 ***
te(roi_dist,angle):mini alpha.local       147.13  208.7   8.857  <2e-16 ***
s(cell_id)                                 14.96   16.0 928.261  <2e-16 ***
```

This indicated that all cell types show significant variation in the smooth surface of RF overlap vs. angle and distance for both local and global stimuli.

Overall, the model explained 62.8% of the deviance.

#### SUPPLEMENTARY FIGURES

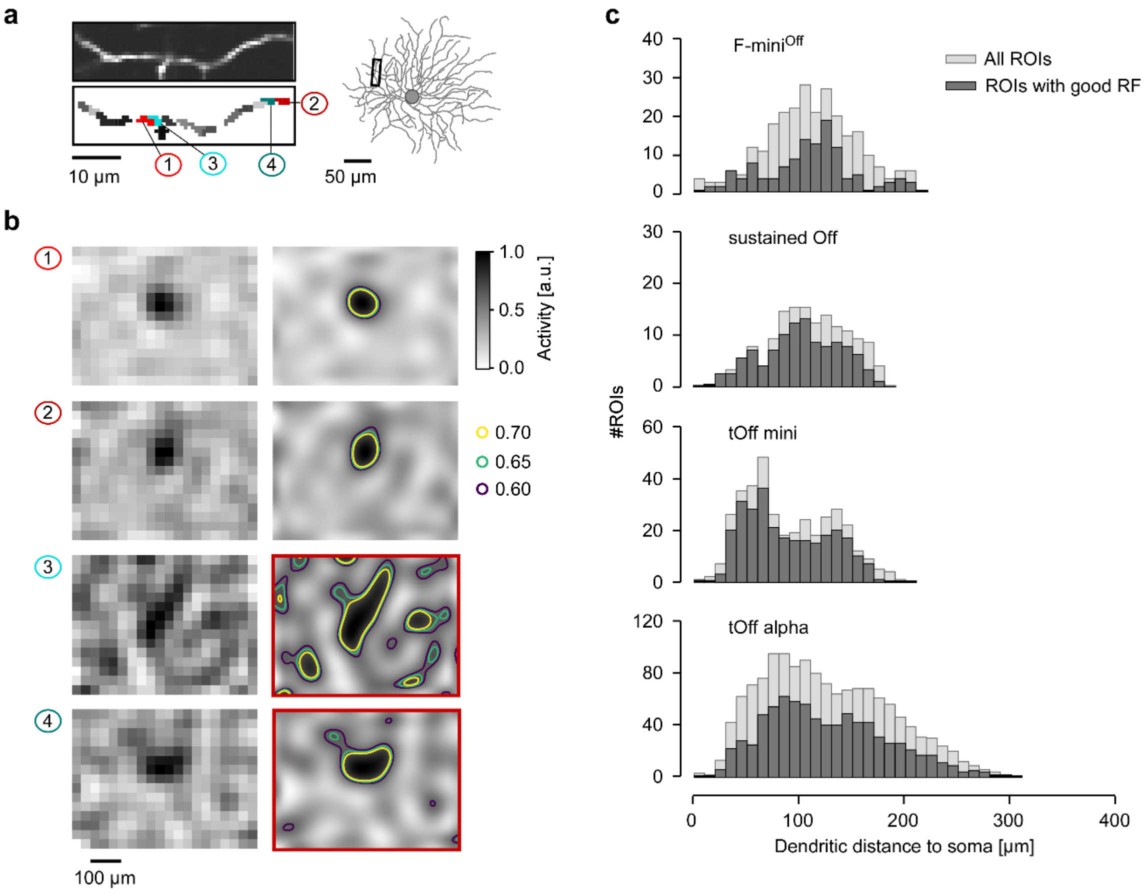


**Supplementary Figure S1 | ROI selection. a**, Exemplary scan field (top left) and automatically generated ROI mask (bottom left) from the region labelled by black rectangle on the reconstructed RGC morphology (right). **b**, Smoothed and normalized RF maps before (left) and after (right) up-sampling for the labelled ROIs indicated in (a). Coloured curves on up-sampled RF maps (right) show RF contours with three different thresholds used for the RF quality test (Methods). Only ROIs that passed the test (a single contour with $Ii < 0.1$, $A_{contour} > 1.8 \cdot 1{,}000$ µm$^2$, at a contour threshold of 0.60; see also Methods) were used for further analysis. Red rectangles around the up-sampled RF maps (right) indicate that RFs did not pass and were discarded. **c,** Histograms of recorded ROIs for tOff alpha (n=17\1,452\850 cells\total ROIs\ROIs that passed the RF quality test), tOff mini (n=5\387\295), sOff (n=4\208\154) and F-mini$^{Off}$ RGCs (n=5\265\126).

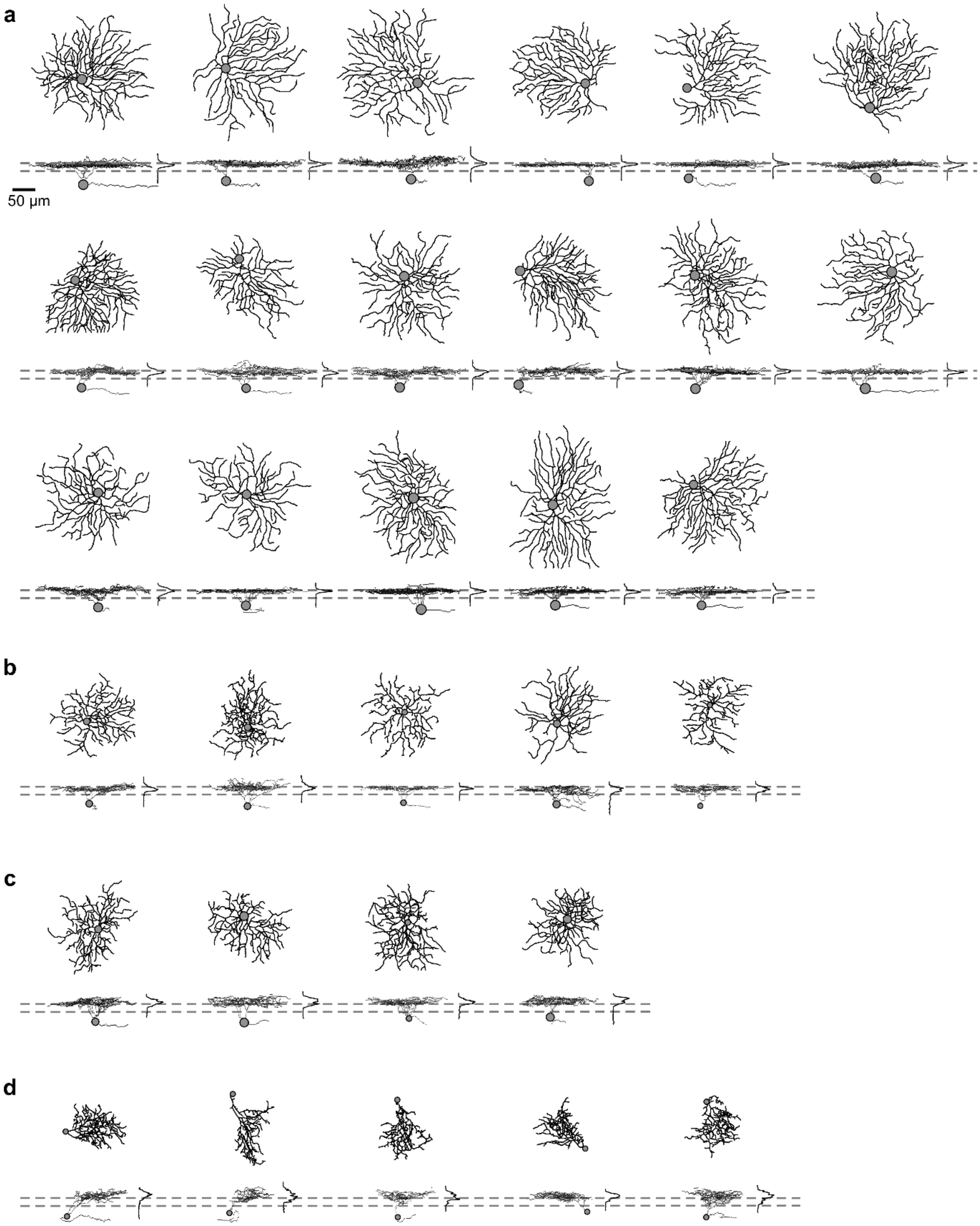


**Supplementary Figure S2 | Morphologies of all recorded RGCs. a-d**, Reconstructed morphologies clustered into RGC types using the algorithm published by (Bae et al., 2018): tOff alpha in (a), tOff mini in (b), sOff in (c) and $\text{F-mini}^{\text{Off}}$ in (d).

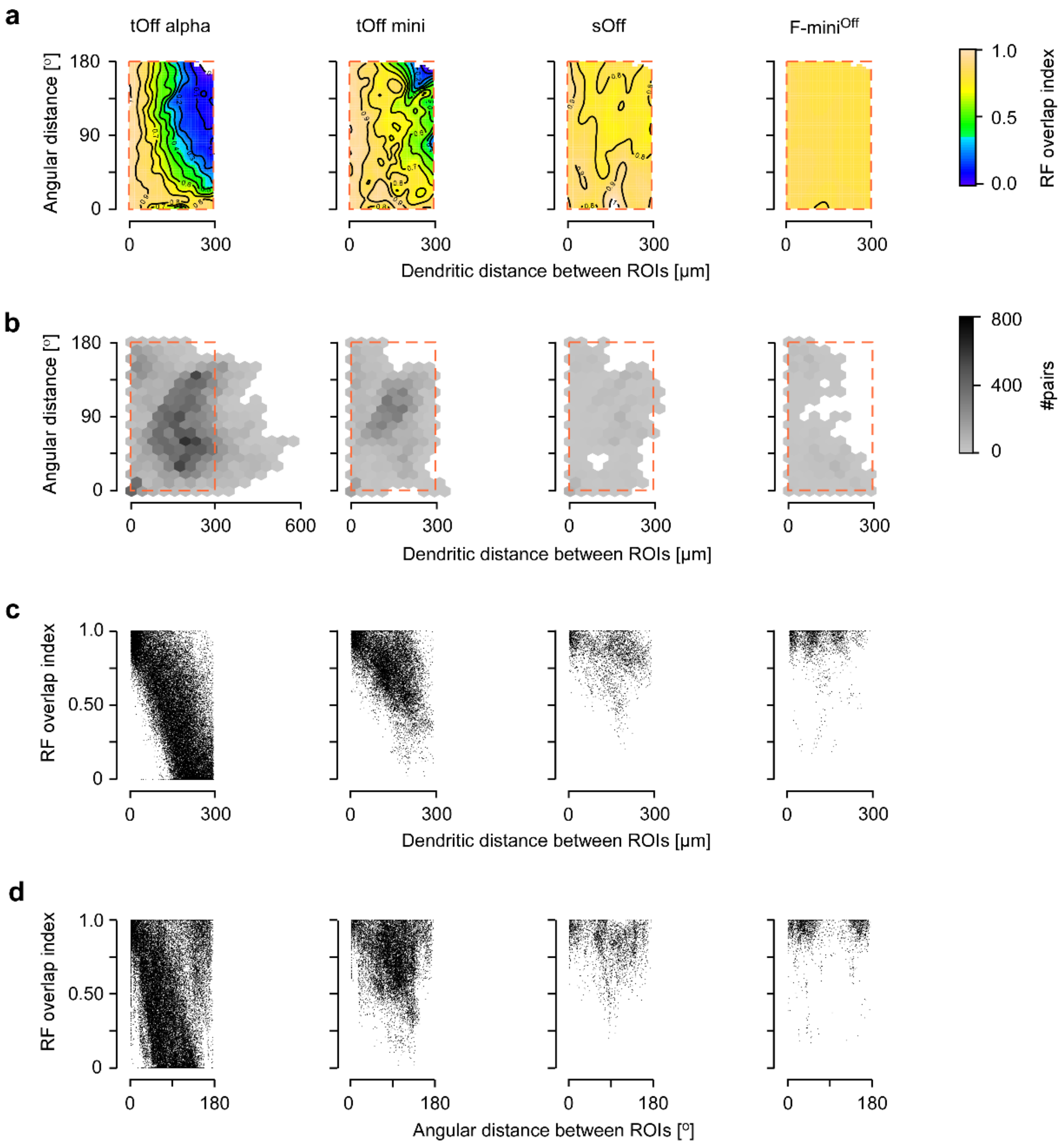


**Supplementary Figure S3 | Dendritic RF overlap data used for statistical comparison. a,** GAM-fitted maps (Methods) of the data shown in main Figure 4c for the plot area marked by dashed red rectangle; colours encode dendritic RF overlap index ($Oi$). **b,** Hexagon maps showing the number of ROI pairs available for estimation of the $Oi$ in tOff alpha (n=17\40,777 cells\ROI pairs), tOff mini (n=5\13,524), sOff (n=4\3,141), and F-mini$^{Off}$ RGCs (n=5\2,097). **c,** RF $Oi$ as a function of dendritic distance between ROIs for the different RGC types, for same plot area as in (a). **d,** Like (c), but with RF $Oi$ as a function of angular distance.

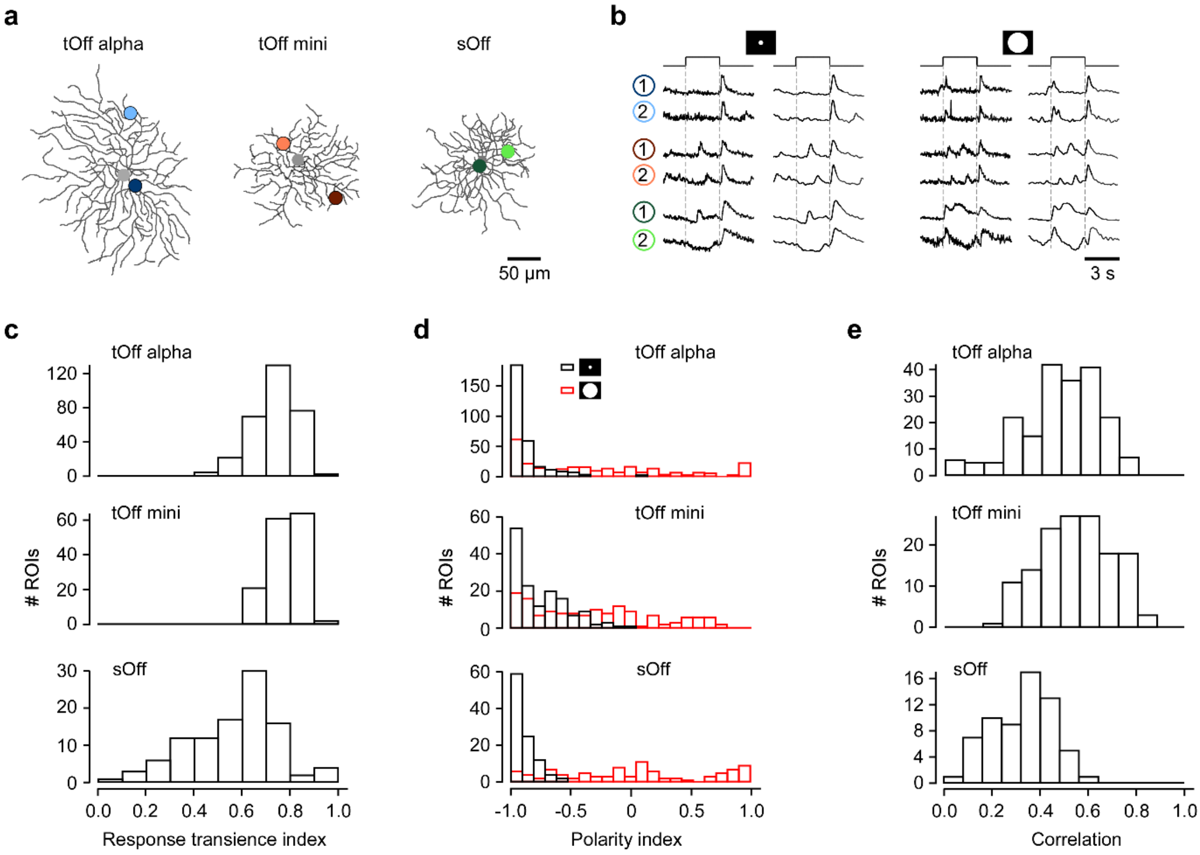


**Supplementary Figure S4 | Temporal features of dendritic responses in different RGC types. a**, Two ROIs overlaid with the reconstructed cells. **b**, Exemplary traces – unsmoothed and binomially smoothed (Methods) – in response to the step section of the local and full-field chirp stimulus recorded from the ROIs in (a). **c**, Histograms of response transience index ($Ti$, Methods) for tOff alpha (n=11\307 cells\ROIs), tOff mini (n=3\148), and sOff RGCs (n=4\103). **d,** Like (c), but with polarity index ($POi$) for tOff alpha (n=11\307\282 cells\ROIs for local\ROIs for full-field chirp), tOff mini (n=3\148\146), and sOff RGCs (n=4\103\93). **e,** Histograms of correlation between local and full-field chirp responses in tOff alpha (n=11\377 cells\ROIs), tOff mini RGCs (n=3\153), and sOff (n=4\129).

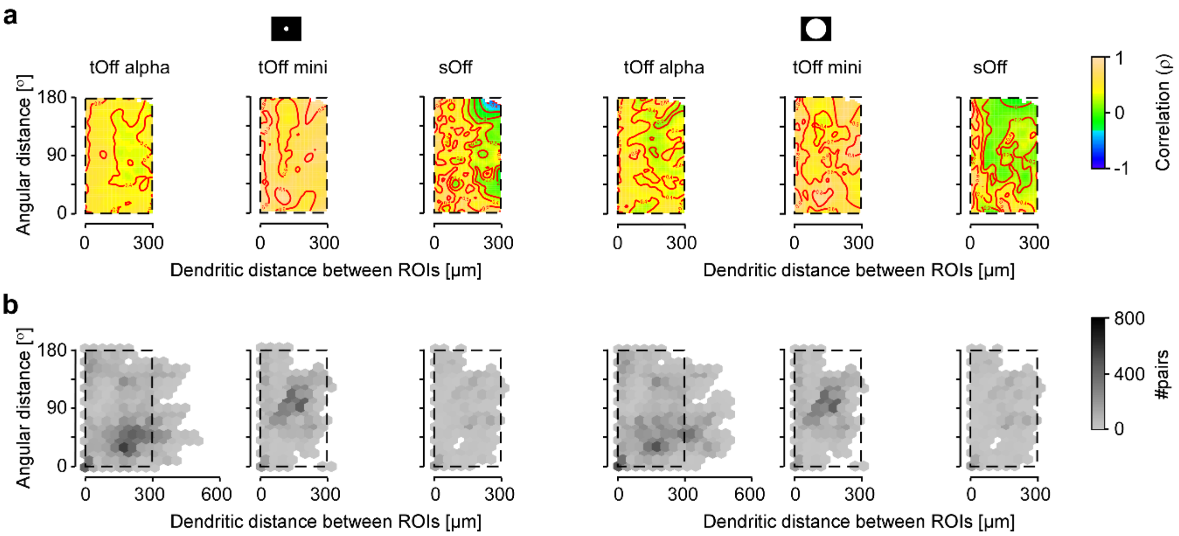


**Supplementary Figure S5 | Local and full-field chirp correlation data used for statistical comparison.** **a,** GAM-fitted maps (Methods) of the correlation data for local (left) and full-field chirp (right) responses shown in main Figure 5d (for plot are marked by dashed black rectangle). **b,** Hexagon maps showing the number of ROI pairs available for estimation of correlation for local (left) and full-field chirp (right) responses.

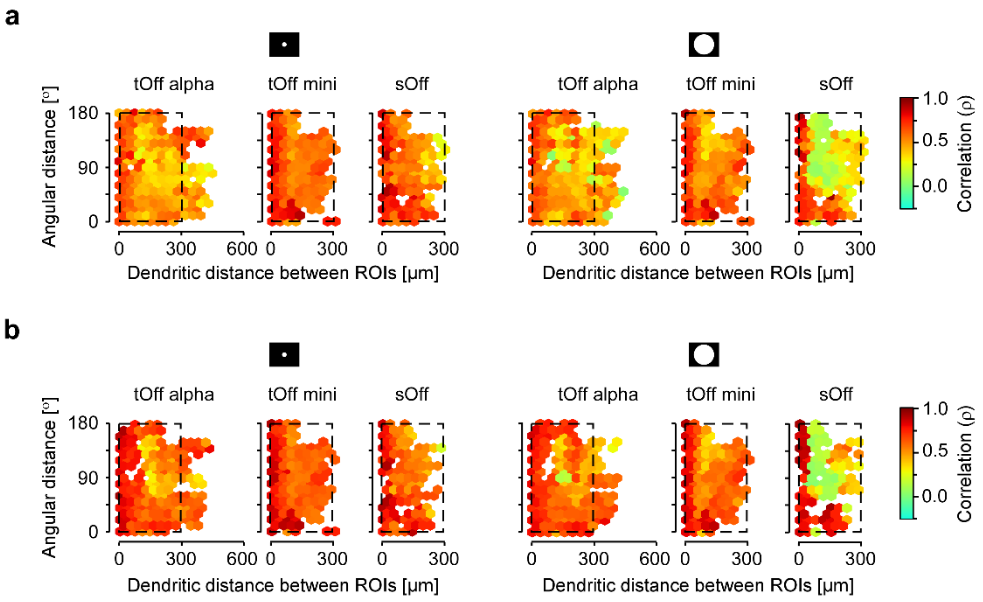


**Supplementary Figure S6 | Temporal correlation across dendrites for different quality thresholds.** **a,** Hexagon maps showing correlation calculated from dendritic signals evoked by local (left) and full-field chirp (right) as a function of angular distance and dendritic distance between ROIs for tOff alpha, tOff mini, and sOff RGCs. Colour encodes correlation; only ROIs with $Qi$ > 0.4 included (Methods). **b,** Same as in (a), but for ROIs with $Qi$ > 0.5. Dashed black rectangles indicate plot area used for statistical analysis (e.g. in main Figure 5d).

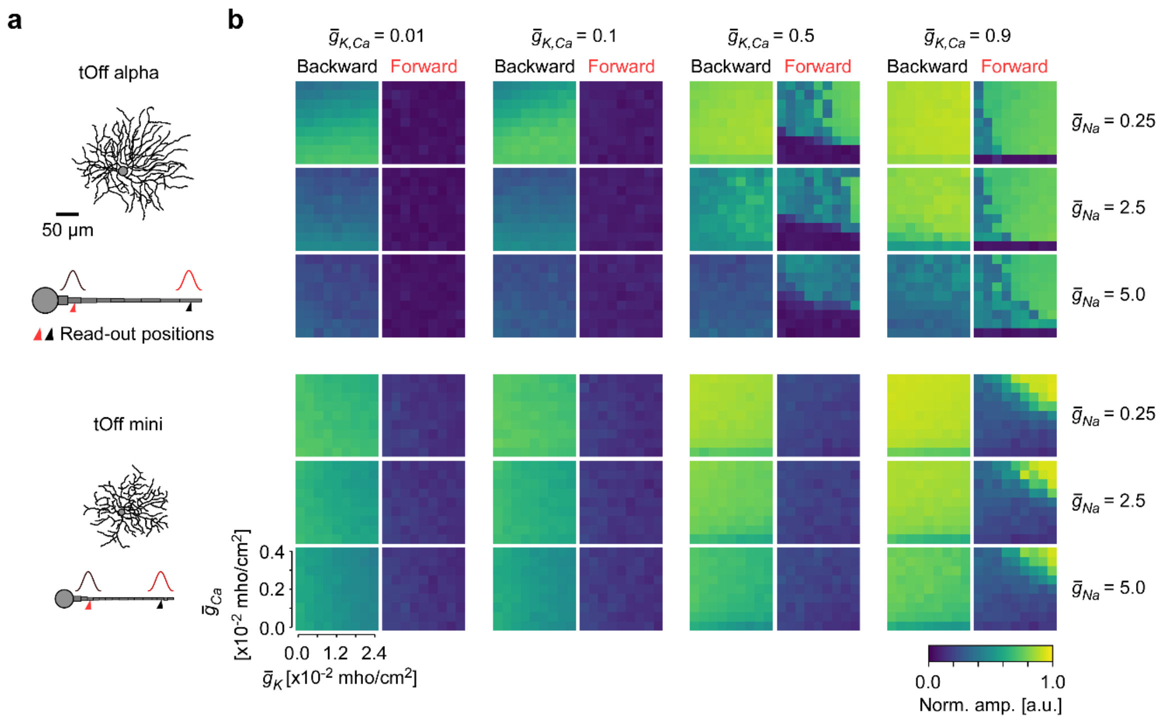


**Supplementary Figure S7 | Simulation of dendritic signal propagation in tOff alpha and tOff mini RGCs. a,** Reconstructed cell morphologies of a tOff alpha and a tOff mini RGC with illustrations of the respective ball-and-stick models (same as in main Figure 6a,c). Simulated inputs at proximal (25 µm to soma) and distal (85% of the total dendrite length to soma) positions indicated as red and black Gaussians, respectively. **b,** Heat maps showing the signal amplitude at the two read-out positions indicated in (c), normalized to the amplitude at the respective input position as a function of ion channel density combinations.

#### SUPPLEMENTARY REFERENCE(S)

Bae, J.A., Mu, S., Kim, J.S., Turner, N.L., Tartavull, I., Kemnitz, N., Jordan, C.S., Norton, A.D., Silversmith, W.M., Prentki, R.*, et al.* (2018). Digital Museum of Retinal Ganglion Cells with Dense Anatomy and Physiology. Cell *173*, 1293-1306 e1219.
